# Genome of an iconic Australian bird: Chromosome-scale assembly and linkage map of the superb fairy-wren (*Malurus cyaneus*)

**DOI:** 10.1101/742965

**Authors:** Joshua V. Peñalba, Yuan Deng, Qi Fang, Leo Joseph, Craig Moritz, Andrew Cockburn

## Abstract

The superb fairy-wren, *Malurus cyaneus*, is one of the most iconic Australian passerine species. This species belongs to an endemic Australasian clade, Meliphagides, which diversified early in the evolution of the oscine passerines. Today, the oscine passerines comprise almost half of all avian species diversity. Despite the rapid increase of available bird genome assemblies, this part of the avian tree has not yet been represented by a high-quality reference. To rectify that, we present the first chromosome-scale genome assembly of a Meliphagides representative: the superb fairy-wren. We combined Illumina shotgun and mate-pair sequences, PacBio long-reads, and a genetic linkage map from an intensively sampled pedigree of a wild population to generate this genome assembly. Of the final assembled 1.07Gb genome, 894Mb (84.8%) was anchored onto 25 chromosomes resulting in a final scaffold N50 of 68.11 Mb. This high-quality bird genome assembly is also one of only a handful which is also accompanied by a genetic map and recombination landscape. In comparison to other pedigree-based bird genetic maps, we find that the zebrafinch (*Taeniopygia*) genetic map more closely resembles the fairy-wren map rather than the map from the more closely-related *Ficedula* flycatcher. Lastly, we also provide a predictive gene and repeat annotation of the genome assembly. This new high quality, annotated genome assembly will be an invaluable resource not only to the superb fairy-wren species and relatives but also broadly across the avian tree by providing a new reference point for comparative genomic analyses.

## Introduction

We present a chromosome-level annotated assembly of the genome of the superb fairy-wren *Malurus cyaneus* (Maluridae). Although genome assembly resources for bird species are accumulating quickly, the phylogenetic coverage of chromosome-scale genomes is currently badly biased (Figure 1). Among the oscine passerines, by far the largest radiation of birds, chromosome level genomes are limited to five species from the Passerides clade. Yet the early evolution of the Oscines involved multiple branching in the Australo-Papuan region in the Oligocene before the emergence of the Corvides and Passerides, the two groups that ultimately gave the oscines their numerical and ecological dominance (Marki et al., 2017; Oliveros et al., 2019). The superb fairy-wren, *Malurus cyaneus*, is an exemplar of the largest clade of that early radiation, the Meliphagides, which has almost 300 species. Furthermore, we complement this high-level assembly with a detailed genetic map from which we can infer the variation in recombination across the genome. Although genome assembly and genetic map resources have been developed separately for various species, there are currently only four other bird species for which we have both resources readily available.

**Figure 1.**
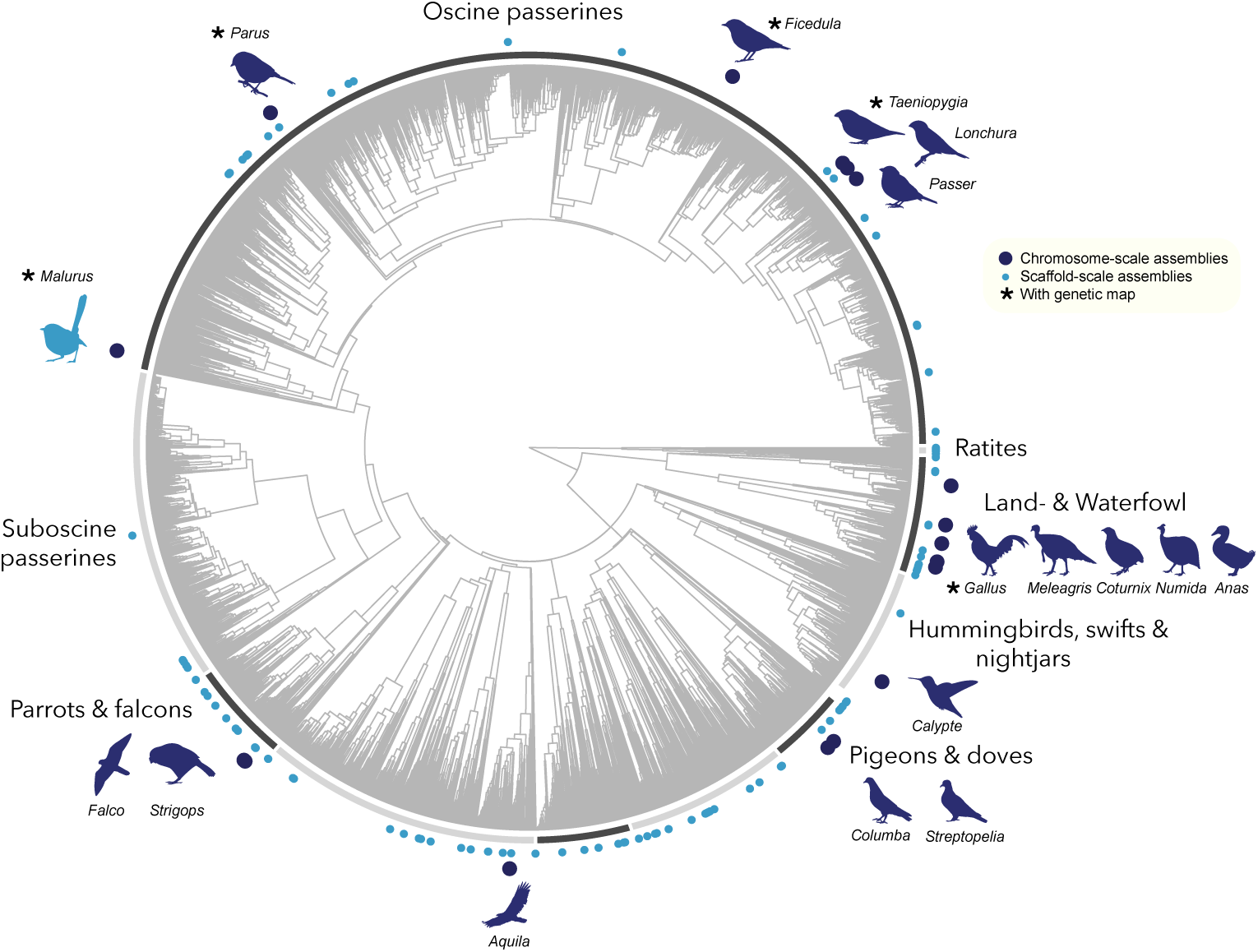
Phylogenetic coverage of available bird genome assemblies. The phylogenetic placement of bird genome assemblies available in NCBI GenBank as of (19 Aug 2019). The dark blue circles and associated silhouettes represent the chromosome-scale genomes and light blue circles represent the scaffold-scale genome assemblies.

Superb fairy-wrens are small, insectivorous birds found throughout south-eastern Australia. The males boast a familiar, striking bright blue breeding plumage contrasted against dark black patches. They have become a model species in behavioural and evolutionary ecology because they combine two conceptually interesting behaviours in an unpredicted way. First, they were the first bird anywhere in the world shown to exhibit cooperative breeding, and the single female on each territory can be assisted by as many as five males in provisioning offspring (Boland & Cockburn, 2002; Rowley & Russell, 1997). Experimental studies provided a textbook example of how limitations to mating opportunities preclude dispersal by young males, which instead defer dispersal and help the breeding pair raise offspring (Pruett-Jones & Lewis, 1990). Second, this cooperative breeding occurs in a novel context. Paternity in this species is dominated by extra-group fertilisations, which are sought by the female during pre-dawn forays to the territories of a small number of attractive males (Cockburn, Brouwer, Double, Margraf, & van de Pol, 2013; Cockburn et al., 2009; Double & Cockburn, 2000; Mulder, Dunn, Cockburn, Lazenby Cohen, & Howell, 1994). As a consequence, males often provision offspring to which they are completely unrelated (Dunn, Cockburn, & Mulder, 1995). This paradox have provoked more than three decades of demographic, behavioural and genetic study of cooperative breeding this species (Cockburn et al., 2016; Cockburn, Sims, et al., 2008). Further issues investigated include mate choice and sexual selection (Cockburn et al., 2016; Cockburn, Sims, et al., 2008; Peters, 2000), inbreeding (Hajduk et al., 2018), dispersal (Cockburn, Osmond, Mulder, Green, & Double, 2003), life history evolution and parental investment (Russell, Langmore, Cockburn, Astheimer, & Kilner, 2007), parent-offspring communication (Colombelli-Négrel et al., 2012), evolution of brood parasitism (Langmore, Hunt, & Kilner, 2003), and behavioural responses to predator risk (Magrath & Bennett, 2012; Magrath, Haff, McLachlan, & Igic, 2015; Potvin, Ratnayake, Radford, & Magrath, 2018), as well as population responses to climate change (Kruuk, Osmond, & Cockburn, 2015; van de Pol, Osmond, & Cockburn, 2012).

Parallel studies of mating systems and extra-group paternity in other *Malurus* species reveal considerable diversity, supporting the idea that evolutionary pathways can be traced through phylogenetically-based comparative analysis (Brouwer et al., 2017; Buchanan & Cockburn, 2013). Other *Malurus* species have also been the focus of studies on breeding biology (Karubian, 2008; Leitão, Hall, Venables, & Mulder, 2019; Varian-Ramos & Webster, 2012), song and vocalizations (Dowling & Webster, 2016; Greig & Pruett-Jones, 2008; Yandell, Hochachka, Pruett-Jones, Webster, & Greig, 2018), plumage and ornamentation (Karubian, 2013; Lindsay, Webster, & Schwabl, 2011), ecology and conservation (Driskell et al., 2011; Murphy, Legge, Heathcote, & Mulder, 2010; Skroblin, Lanfear, Cockburn, & Legge, 2012) and phylogeography (Baldassarre, White, Karubian, & Webster, 2014; Kearns, Joseph, Edwards, & Double, 2009; McLean, Toon, Schmidt, Joseph, & Hughes, 2012). Providing a high-quality, annotated genome assembly sets up the foundation to understand the genetic mechanisms underlying some of the behaviours and natural history traits of the superb fairy-wren and allies.

Upgrading genome assemblies from scaffold-level to chromosome-level provides additional genomic context by orienting genes relative to each other and other genomic features such as centromeres, telomeres, various repeat elements and regulatory regions. Knowledge of this organization aids in understanding how genome architecture can influence variation in evolution within the genome but also provide insight into genome evolution between populations and species (Joseph et al., 2018; Sävilammi et al., 2019; Thind et al., 2018). To date, there are only 16 bird species with genome assemblies classified as “chromosome” level in GenBank (as of 19 Aug 2019). Twelve of these assemblies were assembled *de novo* and four of these were assembled using other reference assemblies as a guide. These assemblies are distinguished from “scaffold” level by higher contiguity where scaffolds are anchored onto chromosomes using either physical (HiC; Burton et al., 2013) or genetic mapping (Fierst, 2015). In the near future, we expect to see an exponential accumulation of bird genome assemblies to be released, particularly from the B10K consortium which is currently sequencing at least one bird species per family (∼300 assemblies; Zhang et al., 2015). Despite this, only a handful of species are in the pipeline to be sequenced to chromosome-level, all of which using HiC chromatin interaction maps (Stiller & Zhang, 2019). Chromosome-scale genomes assembled using a genetic map will remain valuable despite the onslaught of new assemblies being, as yet, limited to a few exemplar species.

The utility of a genetic map extends far beyond upgrading a genome assembly. The maps themselves serve as an invaluable resource by enabling us to associate a particular phenotypic trait to specific loci through QTL mapping (Su et al., 2017), associate different traits through tight linkage of the genes that code for them (Schwander, Libbrecht, & Keller, 2014), and provide an understanding of the variation in genetic diversity within the genome (Burri et al., 2015). The latter has been of particular interest, as of late, as the variation in recombination within the genome provides a hypothesis as to why we might observe particular patterns of variation in genetic diversity, genetic divergence, and gene flow when comparing genomes of populations or species (Comeron Josep M., 2017; Ravinet et al., 2017). It is also of interest to understand how recombination landscape itself varies between individuals, populations, and species and how it may be influenced by selection, demographic history, genomic features, and chromosomal rearrangements (Barton, 1995; Dapper & Payseur, 2017; Ortiz-Barrientos, Engelstädter, & Rieseberg, 2016; Stapley, Feulner, Johnston, Santure, & Smadja, 2017). From the existing chromosome-scale genomes, only four other species have a high-density, SNP-based genetic map to complement the assembly (Backström et al., 2010; Groenen et al., 2009; Kawakami et al., 2014; van Oers et al., 2014). Providing a genetic map and recombination landscape to accompany a high-quality genome assembly greatly expands the array of possible research questions and avenues.

Here we combine both conventional short-read (Illumina shotgun and mate-pair) and long-read sequencing (PacBio) with an extensive pedigree of our long-term study population to achieve a highly contiguous chromosome-level assembly and recombination map. The fairy-wren genome has 894 Mb (out of the 1.07 Gb total assembled) of sequences anchored into 25 chromosomes with a contig and scaffold N50 of 465 Kb and 68.11 Mb, respectively. We also provide comparisons of recombination rate and other genomic features to help understand the sources of variation of these features within the genome. Lastly, we add prediction-based repeat and gene annotations using existing libraries from other bird species. This assembly will greatly facilitate ecological and evolutionary studies of fairy-wrens, and provide a resource for understanding the evolution and diversification of the Meliphagides as a whole.

## Methods

### Reference genome sequencing

We chose a female individual (ANWC:B45704) from the Flinders Island subspecies (*M. cyaneus samueli*) for the reference genome sequencing (Figure 2); a previous microsatellite survey of *M. cyaneus* suggested that this population has the lowest heterozygosity across the species (Etamadmoghadam *et al. undated thesis* unpublished data) making it an ideal candidate for genome assembly. We extracted the DNA from a tissue sample using a standard salting out procedure. For small insert sizes, we prepared two size ranges centered on 250bp and 500bp using the Meyer & Kircher (2010) protocol. The DNA was sheared using a BioRuptor (Diagenode), then we used a double-sided bead size selection to obtain the correct insert size. The 250bp library was sequenced using half a lane of an Illumina HiSeq 2500 (100bp, paired-end) and the 500bp library was sequenced using a lane of Illumina MiSeq (300bp, paired-end). For the three Illumina mate-pair libraries, we prepared insert sizes centered around 3.5Kb, 5.5Kb, and 7.5Kb. Each mate-pair library was sequenced using 1/6th of an Illumina HiSeq 2500 (100bp, paired-end). For the single-cell, long-read libraries, we sequenced across 27 SMRT cells on the Pacific Biosciences RSII platform.

**Figure 2.**
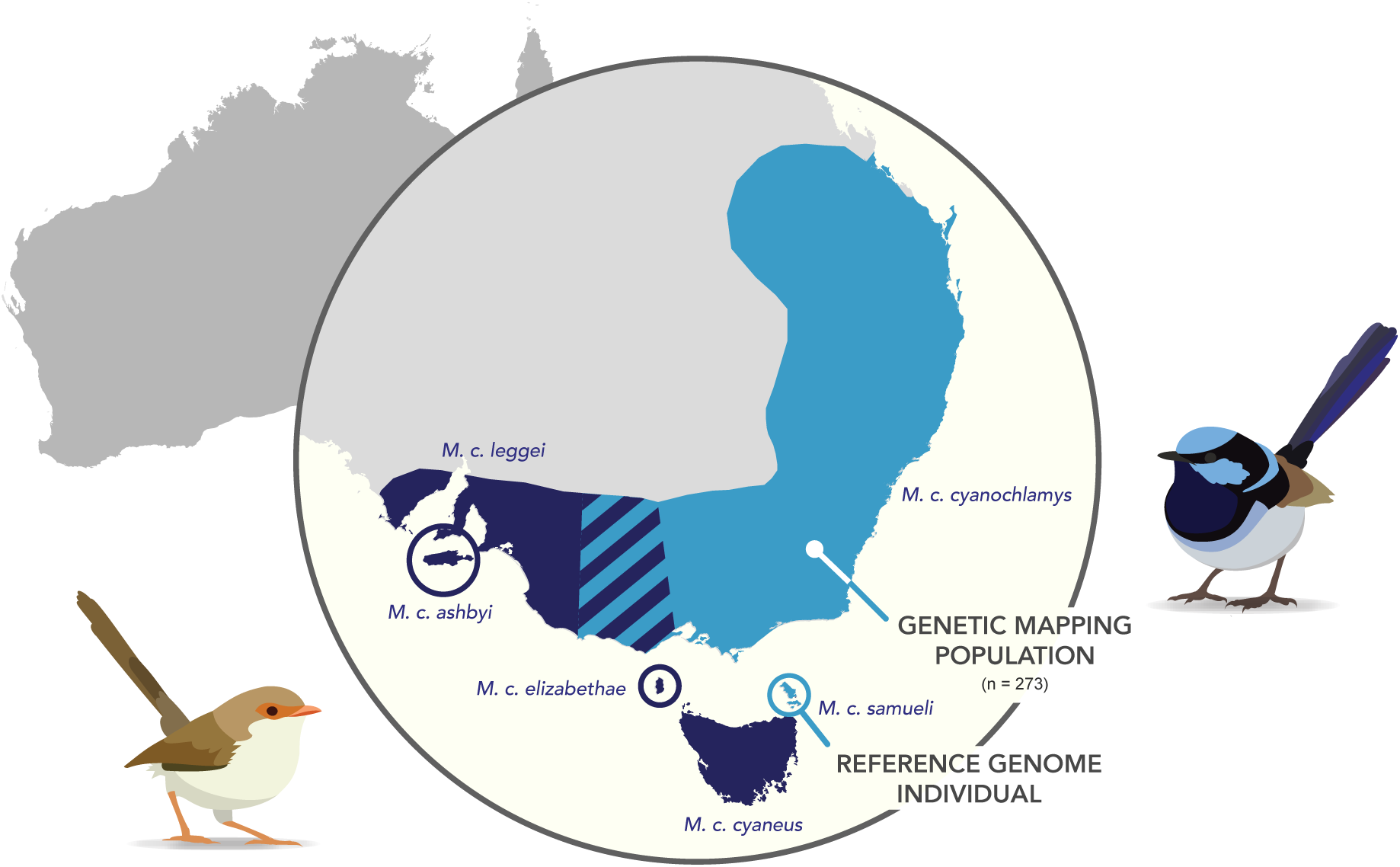
Superb fairy-wren range map and sampling. The entire range of the superb fairy-wren is shown here with the six subspecies designations. The individual chosen for the reference genome (ANWC:B45704) comes from the *samueli* subspecies from Flinders Island next to Tasmania, while the linkage mapping population comes from the mainland *cyanochlamys* subspecies.

### Linkage map sampling and sequencing

We used a subset of the extensively-sampled, wild pedigree of a population of the subspecies *M. cyaneus cyanochlamys* in the Australian National Botanical Garden (Canberra, ACT, Australia) for linkage mapping (Figure 2). Male philopatry and high skew in extra-group mating success resulted in a complex pedigree (Figure S1; Cockburn, Osmond, & Double, 2008; Dunn & Cockburn, 1999)). We chose 273 individuals spanning three to nine generations. Paternity was inferred *a priori* using microsatellite data (and subsequently confirmed using SNP genotyping) and these paternity assignments were used to select the individuals for mapping. We then sequenced tens of thousands of SNPs distributed randomly across the genome using a genome reduction method provided by Diversity Arrays Technology Pty Ltd (Canberra, ACT, Australia; Kilian et al., 2012).

### *De novo* genome assembly

The nuclear genome assembly pipeline can be divided into five main stages which improved contiguity, reduced gaps, and/or improved quality. Each stage is detailed in the supplementary material but will be summarized here (Supplementary methods, Figure S2). The first stage was a short-read (Illumina-only) assembly. The raw Illumina reads for both the short insert sizes and mate-pair libraries were trimmed and cleaned using the *libngs* toolkit (https://github.com/sylvainforet/libngs). The genome size was initially estimated using *SGA preqc* (Simpson, 2014). This size was used to roughly guide the assembly and assessment of its resultant contiguity. We used the ALLPATHs assembler for the initial assembly and scaffolding of the cleaned short-insert and mate-pair Illumina reads (Butler et al., 2008). The second stage was the long-read gap-filling using PacBio subreads. The coverage of the PacBio reads were not sufficient enough to be included in the *de novo* assembly but sufficient for filling gaps. These PacBio reads were first error-corrected using the higher quality Illumina reads using the tool *LoRDEC (*Salmela & Rivals, *2014*). We then used the error-corrected PacBio reads to upgrade the Illumina scaffolds by filling in the gaps and extending the assembly using PBJelly (English et al., 2012). The third stage was anchoring and orienting the scaffolds into chromosomes. This assumes that the genetic order, as inferred from the pedigree-based linkage map, corresponds to the physical order along the chromosome. The details on how the genetic map was built for scaffolding is described in more detail below and in the supplementary methods. We used the software LepMap3 to build our genetic map using the DaRTseq data of our pedigree (Rastas, 2017). Before the scaffolding step, we identified the large misassemblies and split them accordingly. We then used ALLMAPs to anchor and orient the scaffolds using the information from the genetic map (Tang et al., 2015). This stage resulted in new gaps and new associations between scaffolds which led to the fourth stage being another round of gap-filling. Similar to the second stage, the fourth stage used the error-corrected PacBio reads and PBJelly to fill in any new gaps. The fifth and final stage was to polish the genome assembly and convert it to haploid. This stage involved mapping all of the Illumina reads onto the stage four assembly using BWA. We then used Pilon (v.1.22), which uses the information from the alignment to correct incorrectly called bases and small indels (Walker et al., 2014).

The remainder of the steps were curating and naming of the scaffolds. All scaffolds were aligned onto the *Taeniopygia* assembly (v.3.2.4, Zhang, Jarvis, & Gilbert, 2014) using LASTZ and the 25 chromosome-scale scaffolds were named based on the match to the *Taeniopygia guttata* genome, which in turn had been initially named for homology to the *Gallus gallus* genome. The main nomenclature adopted from the *Taeniopygia* genome was the fissions relative to the *Gallus* genome such as chromosome 1A and 4A. W-linked scaffolds were identified using the genotyping of the DArTseq SNPs in the mapping population. Any variants that were found only to be genotyped in females and missing in males were considered putative W-linked SNPs. The scaffolds which these SNPs map to were labeled as unmapped W chromosome scaffolds. The remaining small scaffolds were also aligned to the other passerine chromosome assemblies: *Parus major* (v.1), *Ficedula albicollis* (v.1.5) and *Passer domesticus* (v.1). If these unmapped scaffolds were associated with the same chromosome in two out of the four genomes, they were assigned as ‘unplaced’ to that chromosome. Any remaining scaffolds were labeled as chromosome unknown.

The mitochondrial genome was assembled independently from the nuclear genome. To assemble the mitochondrial genome, we used MITObim which used a reference mitochondrial genome of a closely related species to seed the assembly (Hahn, Bachmann, & Chevreux, 2013). From this initial seed, MITObim iteratively baited and extended the mitochondrial genome assembly until no further extension was recorded. We used a reference assembly from a congeneric species, *Malurus melanocephalus* (GenBank NC024873). We used the Illumina HiSeq (∼250bp insert size) library as it yielded the most consistent results through multiple trials. After choosing the best assembly, we mapped the reads again and corrected any misincorporated sequencing errors using the majority call for each base. Finally, we used MITOS Webserver to identify the location of the genes and tRNAs in the genome (Bernt et al., 2013).

### Genetic linkage mapping for assembly

The third stage of the genome assembly involved linkage mapping to inform the placement and orientation of each scaffold along each chromosome. First, each DaRTseq library was trimmed to remove Illumina adapters and barcode sequences using Trimmomatic (v.0.32, Bolger, Lohse, & Usadel, 2014). The trimmed reads were then mapped onto the PacBio gap-filled scaffolds (stage 2) using BWA mem (Li & Durbin, 2009). The remaining steps follow the pipeline provided by LepMap3 (v. 0.2). This used genotype likelihoods for mapping (Rastas, 2017). The pedigree was split into 37 full-sibling families with a total of 273 individuals. Individuals were included more than once if they belonged to multiple families and grandparents were included to aid phasing.

We first built a framework map by following the LepMap3 (LM3) pipeline. We highlight the relevant parameters here but the detailed description of the mapping can be found in the supplementary methods and Figure S3. The *SeparateChromosomes2* module split the loci into linkage groups which should be associated to a chromosome. We decided on LOD=13 as our cut-off for the linkage groups after testing various cut-offs for generating linkage groups (Figure S4). We then added remaining markers using the *JoinSingles2All* module with a LOD threshold of 10. The *OrderingMarkers2* module finds the best order for markers within a linkage group. This module was run repeatedly and spuriously mapped markers were manually removed. We continued this process until no more spuriously mapped markers remained. The final marker order constituted our ‘framework map’.

Next, we built a ‘forced map’ using the framework map and forcibly grouping all SNPs within a scaffold into the linkage group to which that scaffold belonged. We did another round of *JoinSingles2All* to map any additional scaffolds that were not mapped in the framework map. Within each linkage group, we then filtered out all SNPs that were below a LOD threshold of 3. Finally, we did multiple rounds of the *OrderingMarkers2* and manual curation to build the map we used to anchor and orient the scaffolds.

### Genome assembly assessment and synteny comparisons

We summarized the contiguity of the genome assembly using standard metrics provided by the Assemblathon 2 stats perl script (https://github.com/ucdavis-bioinformatics/assemblathon2-analysis). The contiguity of the genome assemblies were assessed between each major assembly step and the final assembly was compared to other chromosome-level bird assemblies. The completeness and quality of the genome assembly was assessed using BUSCO (3.0.2) analyses which searches the assembly for 4915 universal single-copy orthologs from the Aves (odb9) database (Simão, Waterhouse, Ioannidis, Kriventseva, & Zdobnov, 2015). As for assembly contiguity, this was measured between each assembly step and between the final assembly and existing chromosome-level assemblies. Large-scale synteny between the final fairy-wren genome assembly and existing chromosome-scale assemblies were compared using the LASTZ (v. 1.04.00) alignments.

### Predictive gene and repeat annotation

The predictive gene annotation followed the same annotation pipeline that was run for other avian assemblies in the B10K project for comparability (Zhang et al., 2015). Briefly, the primary gene set was derived from Ensembl85 (from the *Gallus* galGal4 and *Taeniopygia* taeGut2 genome assemblies) totaling 20,194 genes. A supplementary gene set was also compiled using 20,169 human genes, and genes from 71 transcriptomes for a second set. The two rounds of annotation using the two different gene sets involved a rough alignment using tblastn (v. 2.2.2) and generating a gene model using GeneWise (wise2.4.1). We then filtered out short proteins (<30 amino acids), pseudogenes, retrogenes, highly duplicated genes with 70% repeats or with single exon, and redundant or overlapping genes. For *de novo* repeat discovery, we ran RepeatModeler (v. 1.0.8) on the assembly to create a *Malurus* specific repeat library. To characterize the final repeat content, we ran RepeatMasker (v. 4.0.7) using the database containing the *Malurus* specific repeat library and those from other avian databases: *Taeniopygia guttata, Ficedula albicollis,* and *Corvus cornix (Vijay et al., 2016)*.

### Recombination landscape and comparisons

The process to generate the recombination landscape was identical to the linkage mapping but used the information from the physical position on the final genome assembly as the physical order of the markers. First, we remapped the DaRTseq reads on the final polished genome assembly using BWA. The first stages of the LM3 pipeline were run as previously. As with the forced map, within each chromosome we filtered out markers which fell below the LOD score limit of 3. We then ran the *OrderMarkers2* module using the physical order and obtained the sex-specific and sex-averaged genetic maps for the 24 autosomes and the male-specific map for the Z chromosome.

We obtained recombination rate estimates from the Marey map representation: genetic distance plotted against physical distance. First, we smoothed the map using a LOESS (local polynomial regression) smoothing with a span of 0.2. This regression smooths over windows of a fixed number of SNPs instead of physical length to reduce bias in regions where more SNPs were recovered. Using this regression, we estimated recombination rates in 200kb nonoverlapping windows and correlated these estimates with other genomic features. GC content was calculated as the percent of G or C bases within the given window. The gene density was the percentage of coding sequence within the 200kb window. Relative distance from chromosome end was standardized from 0 to 1, 0 corresponding to the chromosome end and 1 corresponding to chromosome center. Prior to analyses, recombination rate was log base 10 transformed to reduce skewness. Gene density and GC content were both square root transformed. We explicitly tested the correlation of recombination rate to both gene density and distance to chromosome end using Pearson’s R. GC content was not explicitly correlated with recombination rate as this feature was not an explanatory variable but rather likely a byproduct of the variation in recombination (Bolívar, Mugal, Nater, & Ellegren, 2016; Clément & Arndt, 2013). To minimize the effects of spatial autocorrelation within the genome, we performed permutation tests to obtain the distribution of the Pearson’s R statistic under a null model (shuffled recombination rate and explanatory variables among windows) and our observed data. Furthermore, we performed bootstrapping by subsampling 20% of the windows with replacement. We performed 2000 iterations of bootstrapping to obtain the null distribution of Pearson’s R and another 2000 for the observed distribution. We included a separate representation of the effect of the distance to chromosome end by comparing the log base 10 transformed recombination rate to the log base 10 transformed exact distance to chromosome end in megabases. We then performed a LOESS smoothing of this correlation to illustrate the shift in recombination rates from the ends of the chromosomes to the center. Between chromosomes, we also compared recombination rate (genetic map distance / chromosome length) to chromosome length and gene density (number of genes / chromosome length). Additionally, we qualitatively compared the fairy-wren recombination landscape with the pedigree-derived recombination landscapes of *Gallus gallus (Groenen et al., 2009) Taeniopygia guttata* (Backström et al., 2010) and *Ficedula albicollis* (Kawakami et al., 2014). The *Parus* genetic map was not included as the analyzed data were readily available.

## Results

### Genome sequencing effort

The raw data comprised 128,683,241 read pairs from the 250bp insert library, 12,456,651 read pairs from the 500bp insert library, 47,009,583 from the 3.5kb mate-pair library, 56,672,592 from the 5.5kb mate-pair library, and 39,069,768 from the 7.5kb mate-pair library. The SGA preqc results estimated the genome size to be 1.07 Gb and 1.04 Gb from the 250bp and 500bp insert size libraries, respectively. Using the 1.07 Gb genome size estimate, the sequencing coverage corresponded to 24.0x from the 250bp insert library, 7.0x from the 500bp insert library, 8.7x from the 3.5kb mate-pair library, 10.5x from the 5.5kb mate-pair library, and 7.3x from th 7.5kb mate-pair library for a total of 55.7x coverage from Illumina sequencing. After filtering and cleaning of the raw reads, we retained 91,417,772 read pairs and 21,503,225 merged or unpaired reads from the 250bp insert size Illumina library and 6,303,745 read pairs and 3,496,087 merged or unpaired reads from the 500bp insert size Illumina library. For the mate pair libraries, we retained 12,287,646 read pairs for the 3.5kb insert size, 15,656,984 read pairs for the 5.5kb insert size and 9,326,889 for the 7.5kb insert size libraries (Table S1).

From the 27 PacBio RSII SMRT cells, we sequenced 2,676,850 subreads. The mean length of the subreads was 8.7kb with the longest subread being 53.6kb. The total coverage of the PacBio filtered subreads was 22x for an estimated 1.07Gb size genome. The distribution of subread lengths can be found in supplemental figure S5. Of the total number of reads, 1,271,158 (47%) was cleaned up by LoRDEC.

### Genome assembly

Here we will focus on the stages of genome assembly but the specific sections that require more attention (such as linkage mapping and misassembly detection) will be discussed in detail below. Stage 1 yielded a scaffold N50 of 6.0Mb and a contig N50 of 15kb. The size of the scaffold assembly was 1.01Gb (94.4% of the predicted 1.07Gb genome size) with a total gap length of 113Mb. Stage 2 yielded a marginally increased scaffold N50 of 8.0Mb and a substantial 31-fold increase in contig N50 to 465kb. Additionally, the gap length was greatly reduced to 27Mb. The size of the scaffold assembly was 1.05Gb (98.1%) after stage 2. The largest contribution of the PacBio reads here was the increase in contig N50 and reduction of the gap length. Stage 3 used the genetic map to place and orient the scaffolds into chromosomes. Since our subsampling of the pedigree only consisted of 273 individuals, we were only able to assembly 25 chromosomes with confidence and not all are as large as their homologs in other bird species. This step anchored 235 scaffolds (894 Mb = 84.8% of the genome) into chromosomes and oriented 198 scaffolds (830Mb = 78.8% of the genome). This left 160Mb (15.2%) of the genome unplaced. The scaffold N50 increased to 67.7Mb (∼8.5-fold increase) but the contig N50 stayed approximately the same (465kb). The total scaffold assembly length is also at 1.05Gb. The gap length remains at 27Mb. Stage 4 yielded only a marginal increase in scaffold N50 to 68.1Mb with an increase in the contig N50 to 540kb. This stage elevated the assembly length to the predicted length of 1.07Gb. This further reduced the gap length to 16Mb. The final stage (5), had a final scaffold N50 of 68.11Mb and contig N50 of 560kb. The total assembly length of the scaffold was 1.07Gb and the contig was 1.06Gb, both of which are around the estimated genome length. In total there were 4,329 scaffolds and 15,027 contigs. The final gap length remained at 16Mb. The BUSCO single-copy gene search found 89.4% complete single copy genes (out of 4915 genes) from the stage 1 assembly and found 92.2% complete single copy genes from stage 5 assembly. The progression of contiguity and the BUSCO assessment of the stages of genome assembly can be found in Figure 3. The assembled chromosome sizes can be found in Table 1. The final MITObim mitochondrial genome assembly size was 17,031bp. The annotation resulted in the expected 13 genes (COI-III, ND1-6, cytB, ATP6 and ATP8), 2 ribosomal RNAs (12S and 16S), and 22 tRNAs (Figure S6).

**Figure 3.**
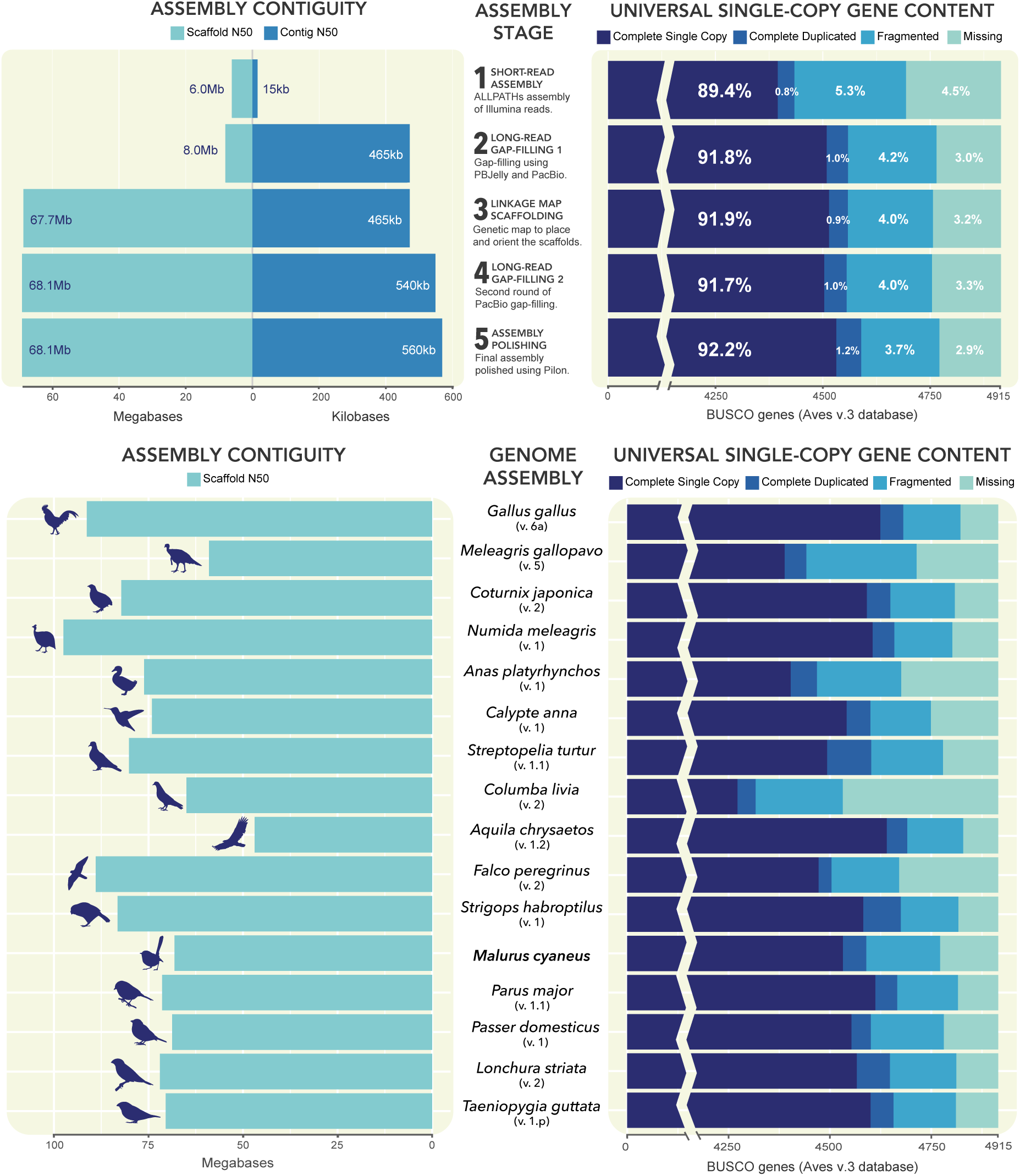
Genome assembly quality assessments. The assembly contiguity metrics on the left plots show scaffold and contig N50 estimated and the assembly completeness metrics on the right plots show the BUSCO gene search using the aves database. The plots on the top row show the progression of increasing assembly quality between the five stages of genome assembly. The plots on the bottom row compare assembly quality between bird genomes, showing the *Malurus* assembly to be comparable to other chromosome assemblies.

**Table 1.** Summary chromosome metrics. Table summarizing the assembly lengths, linkage map lengths, and annotation of each chromosome in the final superb fairy-wren genome assembly.

| Chrom. | Size (Mb) |  | GC% | Genetic Distance (cM) |  |  | Genes<br>(unplaced) |
| --- | --- | --- | --- | --- | --- | --- | --- |
|  | (unplaced) |  |  | Average | Male | Female |  |
| Chrom 1 | 125.32 | (1.70) | 39.96 | 90.11 | 95.55 | 85.58 | 1307 (7) |
| Chrom 1A | 68.11 | (1.56) | 39.92 | 74 | 84.65 | 63.91 | 846 (8) |
| Chrom 2 | 163.99 | (0.52) | 40.06 | 104.54 | 115.68 | 94.83 | 1788 (4) |
| Chrom 3 | 108.86 | (0.67) | 39.83 | 75.75 | 92.09 | 60.07 | 1111 (15) |
| Chrom 4 | 72.11 | (0.60) | 39.48 | 94.59 | 99.7 | 90.28 | 777 (1) |
| Chrom 4A | 18.69 | (1.28) | 44.01 | 48.56 | 52.5 | 45.16 | 303 (54) |
| Chrom 5 | 54.65 | (5.10) | 41.68 | 86.62 | 84.02 | 90.64 | 907 (137) |
| Chrom 6 | 33.73 | (1.01) | 41.96 | 66.35 | 73.08 | 60.36 | 555 (16) |
| Chrom 7 | 35.83 | (1.65) | 41.13 | 84.07 | 89.82 | 79.96 | 509 (51) |
| Chrom 8 | 30.90 | (0.20) | 42.47 | 84.7 | 90.56 | 80.01 | 532 (3) |
| Chrom 9 | 24.52 | (0.04) | 43.31 | 54.55 | 56.75 | 52.83 | 439 (0) |
| Chrom 10 | 20.32 | (0.00) | 43.56 | 78.61 | 82.08 | 75.93 | 416 (0) |
| Chrom 11 | 19.37 | (0.22) | 42.46 | 47.16 | 52.09 | 42.8 | 303 (14) |
| Chrom 12 | 21.60 | (0.04) | 44.12 | 60.19 | 62.79 | 58.47 | 364 (0) |
| Chrom 13 | 19.04 | (0.02) | 45.45 | 46.88 | 43.06 | 51.09 | 376 (0) |
| Chrom 14 | 19.34 | (0.02) | 45.38 | 79.74 | 75.23 | 86.83 | 490 (0) |
| Chrom 15 | 14.69 | (0.13) | 46.33 | 67.23 | 61.94 | 73.41 | 375 (6) |
| Chrom 17 | 12.35 | (0.01) | 48.20 | 50.47 | 45.97 | 55.39 | 294 (0) |
| Chrom 18 | 12.16 | (1.30) | 46.76 | 48.22 | 52.08 | 44.82 | 308 (36) |
| Chrom 20 | 14.96 | (0.06) | 46.77 | 47.28 | 44.82 | 50.17 | 341 (0) |
| Chrom 21 | 5.21 | (2.14) | 47.45 | 25.93 | 26.52 | 25.75 | 154 (77) |
| Chrom 23 | 3.93 | (1.45) | 48.84 | 26.9 | 30.21 | 24 | 96 (43) |
| Chrom 27 | 2.33 | (2.18) | 50.26 | 17.14 | 19.27 | 15.25 | 138 (120) |
| Chrom 28 | 3.94 | (2.20) | 50.74 | 22.16 | 22.92 | 21.6 | 173 (67) |
| Chrom W | - | (0.39) | - | - | - | - | - (3) |
| Chrom Z | 68.91 | (16.80) | 37.23 | - | 72.81 | - | 581 (112) |
| Unknown | - | (62.72) | - | - | - | - | - (1202) |
| TOTAL | 1078.87 |  | 42.04 | 1618.3 | 1686.92 | 1583.19 | 15458 |

### Genetic linkage mapping for chromosome level assembly (Stage 2 to Stage 3)

We recovered 35,276 SNPs after mapping the DArTseq reads onto the stage 2 assembly. The *SeparateChromosomes2* analysis with the LODlimit of 13 yielded 150 linkage groups with 7331 markers (minimum 3 markers per linkage group). From the linkage groups, 36 had at least six markers or more. For the remaining stages we retained 36 linkage groups (6589 loci) ensuring each group had multiple markers that mapped to two or more scaffolds. After joining additional markers to these 36 groups, we retained a total of 11,414 markers. Multiple iterations of the *OrderMarkers2* module and manual curation eventually resulted in 26 linkage groups that we can confidently assign to chromosomes. Some of the smaller linkage groups were associated to the same microchromosome and were merged when appropriate. Since average recombination rates were markedly higher in microchromosomes, they would have likely require a lower LOD score limit relative to the macrochromosomes. The first stage of the forced map resulted in 32,708 SNPs associated to an existing linkage group. Adding new scaffolds onto the existing map resulted in an additional 390 SNPs being mapped. The final *OrderingMarkers2* iterations and manual curation dropped one linkage group which was nested within a single scaffold. This was the final set of 25,078 SNPs in 25 linkage groups that was used for anchoring and ordering of the scaffolds.

The linkage mapping consistently associated markers between two pairs of chromosomes suggesting potential fusions relative to the *Taeniopygia* karyotype. Markers associated with *Taeniopygia* chromosome 26 were consistently associated with chromosome 5 and markers from chromosome 19 were consistently associated with chromosome 2. This is supported by karyological evidence which showed a diploid number of 72 for the *Malurus* and 80 for the *Taeniopygia* (Les Christidis *pers. comm*). Since there were not enough data to reconstruct all of the microchromosomes, we were unable to fully characterize the chromosomal rearrangements with confidence. A summary of the number of SNPs retained for each step can be found in supplementary figure S3.

### Gene and repeat annotation

In total, the gene annotation pipeline predicted 15,458 genes across the genome. Of these, 13,483 (87.2%) were annotated in assembled chromosomes, 774 (5.0%) were annotated in unplaced scaffolds associated with chromosomes, and 1,202 (7.8%) were annotated in scaffolds of unknown location. Of the unplaced scaffolds, we annotated 3 genes associated with the W chromosome (Table 1). With the repeat annotation and masking, RepeatMasker masked 7.90% of the total genome length. Of this fraction of the genome, 41.1% were LINEs, 28.5% were LTR elements, 1.4% were SINEs, 1.0% were DNA elements, and 8.2% were unclassified. Of the remaining, 15.4% were simple repeats, 3.8% were low complexity, 0.6% were small RNAs, and 0.5% were satellite DNA (Table S2).

### Comparison with existing chromosome-scale assemblies

We compared the final *Malurus* genome assembly to 16 other chromosome-scale bird genome assemblies for contiguity, completeness, and synteny. The scaffold N50 of the other genome assemblies ranged from 46.93 to 97.48 (mean=74.7 Mb ± 12.61 Mb s.d.) while the *Malurus* was at 68.1 Mb, and hence falls within the contiguity range of other chromosome-scale bird genome assemblies. The same BUSCO search was run for the other bird genomes. The *Ficedula* genome was omitted due to an undeterminable error during the gene search although the other genomes were sufficient for comparison. On average, 92.15% (± 0.02%) of the 4915 aves (odb9) genes were recovered as complete and single-copy for the other genomes and 92.2% for the superb fairy-wren assembly (Figure 3).

Synteny of the macrochromosomes is largely conserved between the fairy-wren and other bird genome assemblies (Figure 4). While most assemblies had chromosomes which were named after the homology to the *Gallus gallus* genome, the *Meleagris gallopavo*, *Numida meleagris* and *Strigops habroptilus* chromosomes were named from decreasing size. Despite the crossing of lines in the Circos plots being largely due to naming convention, there are still some notable chromosomal rearrangements in these other genomes. The *Falco peregrinus* genome, like other falconid genomes, have previously been shown to have many rearrangements (Nishida et al., 2008; O’Connor et al., 2018) and this is still true compared to *Malurus*. Interestingly the other raptor genome, *Aquila chrysaetos*, also exhibits high levels of rearrangements. Although both *Falco* and *Aquila* have raptorial lifestyles, they belong to different and distantly related clades (Falconiformes and Accipitriformes, respectively). In the species sampled here, *Strigops* is the closest relative of *Falco* (Figure 1). The Circos plots also show likely fusions or translocations of microchromosomes to macrochromosomes in the *Malurus* genome relative to all other genomes. This may be true as evidenced by fewer chromosomes in the *Malurus cyaneus* karyotype but can also be due to misassemblies. Currently, there are not enough data to distinguish these alternatives.

**Figure 4.**
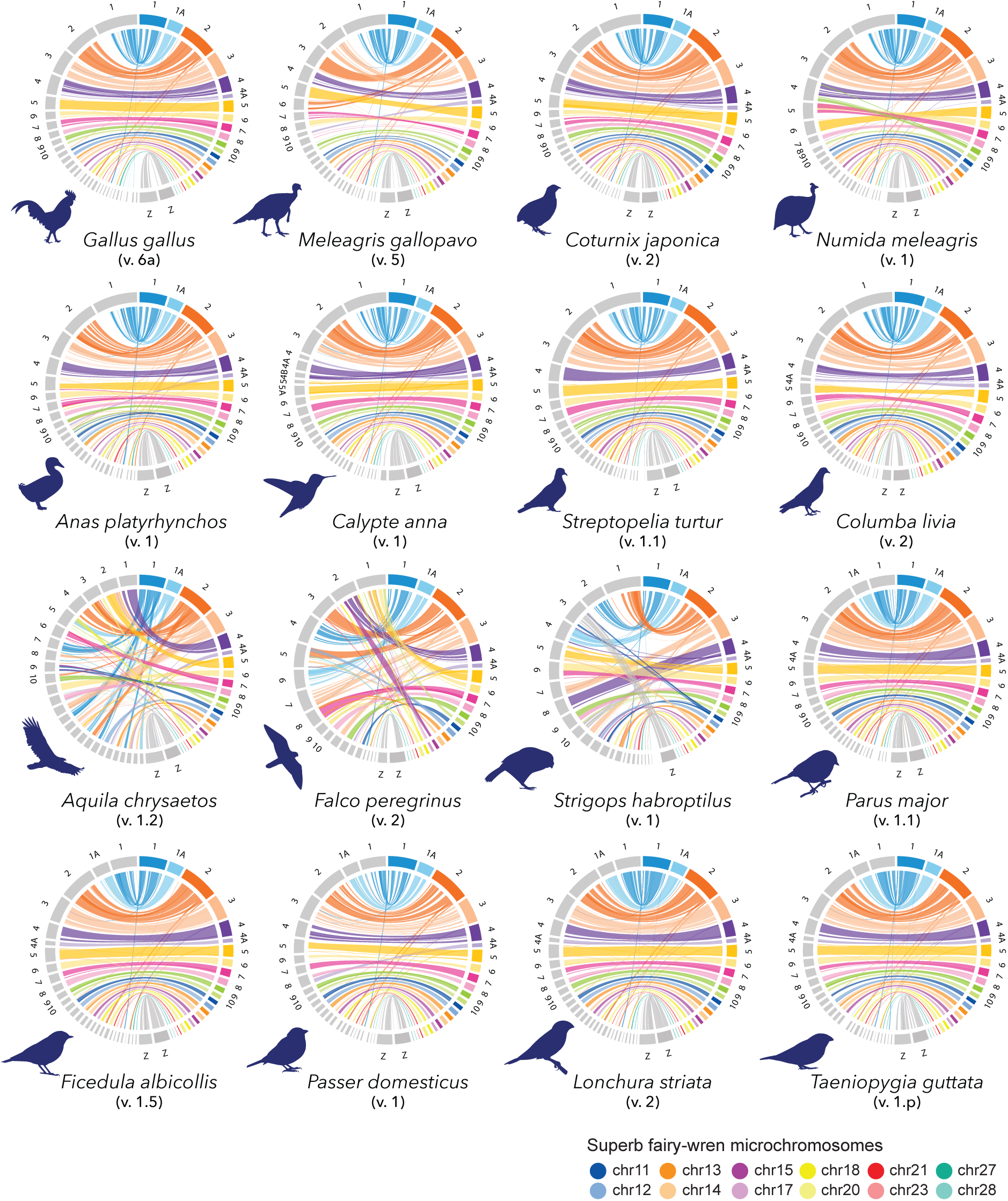
Superb fairy-wren synteny plots. These plots show pairwise synteny comparisons between the *Malurus* chromosomes and the chromosomes of 16 other bird assemblies.

### Genetic map and recombination landscape comparisons

On average, the male-specific, autosomal genetic map of the fairy-wren was longer than that of the female-specific map (Figure S7). This resulted in the total male-specific map being 1.08 times longer than that of the females. Other passerine systems also show deviation from sex equal recombination rates but differ regarding which sex has the longer map. In the *Ficedula albicollis* system, the male genetic map is 1.13 longer than that of the females and in two *Parus major* populations, the female maps are longer by a factor of 1.04 and 1.05 (Kawakami et al., 2014; van Oers et al., 2014). The difference between male and female map lengths also varied with respect to chromosome (Table 1). When map length is converted to recombination rate within the chromosomes, it was not always consistent that male-specific local recombination rate was higher than that of the female-specific rate.

We compared the sex-averaged genetic distance (cM) along the length of the chromosome between different species: *Malurus cyaneus*, *Gallus gallus*, *Taeniopygia guttata* and *Ficedula albicollis* genetic maps (Figure 5). While the detailed comparison of the *Ficedula* map has emphasized that their map is more similar to the *Gallus* than that of the *Taeniopygia*, the *Malurus* map is more concordant with the *Taeniopygia* map (Kawakami et al., 2014), despite the closer phylogenetic distance between *Ficedula* and *Taeniopygia*. The *Malurus* and the *Taeniopygia* maps were both shorter in comparison to the other two maps. Even in chromosome 2, where the *Taeniopygia* genetic map has usually been omitted due to inconsistencies, the *Malurus* map was shorter than that of the other two species suggesting generally lower recombination rates. This pattern was most apparent in the largest chromosomes. Within the smaller chromosomes, the patterns were less consistent where the *Malurus* genetic map would either be more consistent with the *Gallus* and *Ficedula*, more consistent with the *Taeniopygia*, or intermediate. In having recombination deserts across most macrochromosomes, the *Malurus* map is more similar to the distantly related *Taeniopygia* than the more closely related *Ficedula*.

**Figure 5.**
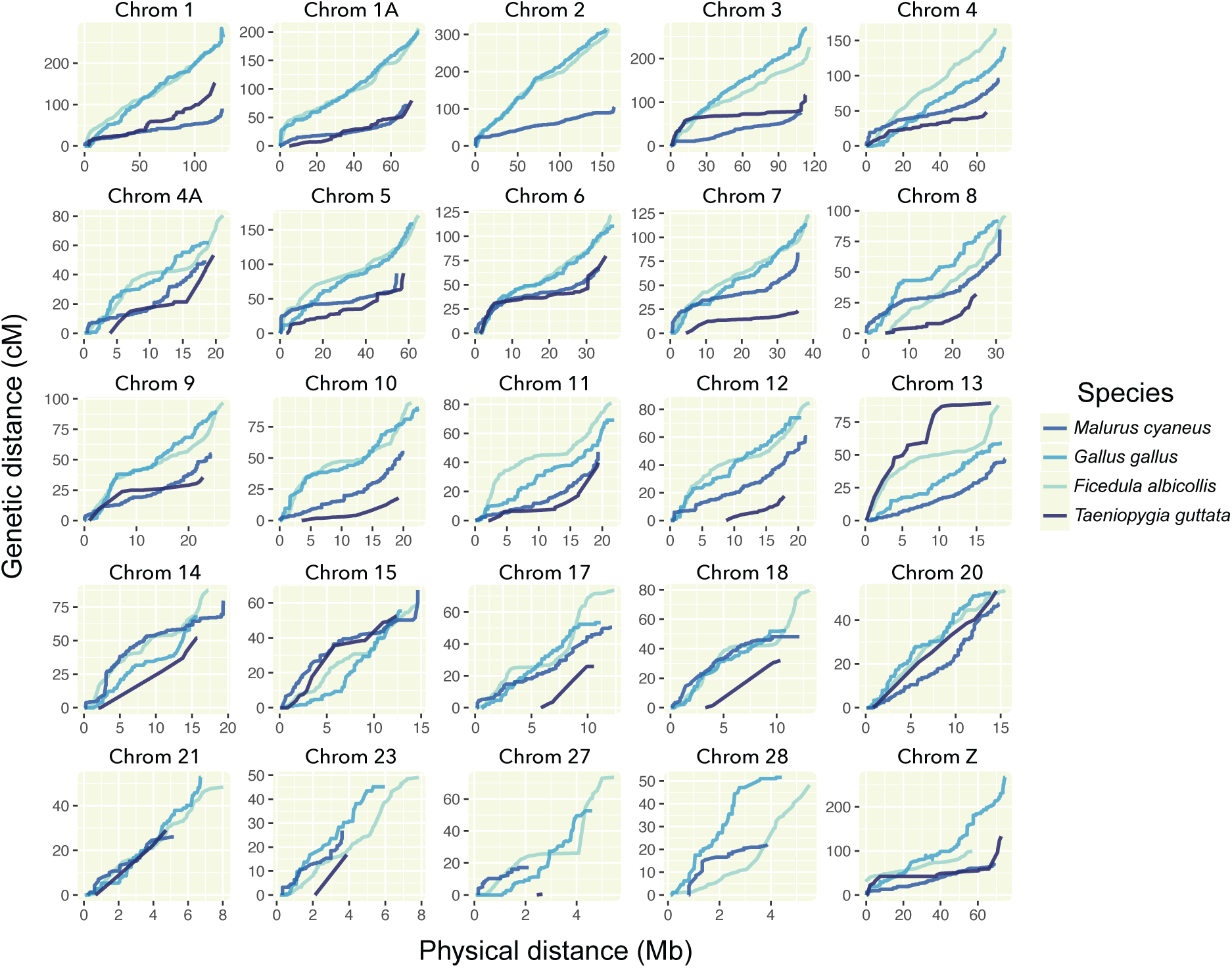
Comparative Marey maps. Sex-averaged genetic distances (cM) plotted against physical distances (Mb) across 25 autosomes and the Z chromosome. The pedigree-based linkage maps of the chicken (*Gallus gallus*), collard flycatcher (*Ficedula albicollis*), and zebrafinch (*Taeniopygia guttata*) are plotted with the *Malurus* map for comparison. The remaining *Malurus* autosomes were omitted due to short size and lack of information.

We compared the recombination landscape (cM/Mb) across the genome estimated from the genetic map with various genomic features (Figure 6). All comparisons were made using measures in 200kb non-overlapping sliding windows. The gene density had the lower strength of association (Pearson’s R = 0.12 ± 0.03) but was still significantly different from zero and null distribution. This positive correlation between gene density and recombination rate was consistent within and between chromosome comparisons. The macrochromosomes were the least gene-dense and had the lowest average recombination and the microchromosomes were both gene-dense and had high average recombination rates. The higher correlation from our comparisons was the relative distance from chromosome end (Pearson’s R = −0.59 ± 0.02). The negative relationship showed decreasing recombination rate from the end to the center of the chromosome. The LOESS smoothed relationship between recombination rate and distance from chromosome end also showed a rapid decrease in recombination rate (Figure 6). This can also be observed in the recombination rate plotted across the length of the chromosomes (Figure S8).

**Figure 6.**
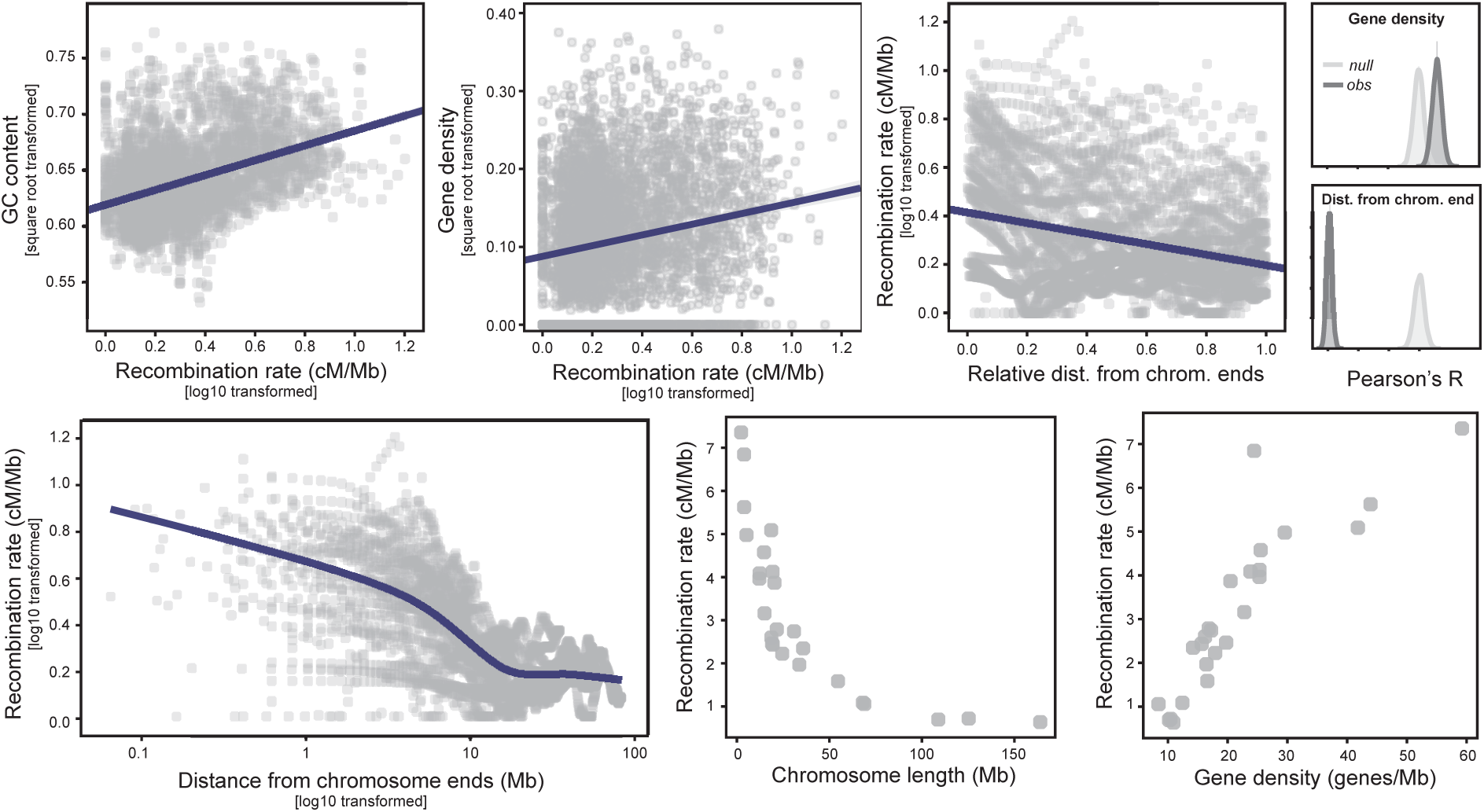
Recombination rate correlations. Plots showing relationship between recombination rate and different genome features. From left to right the top panel shows the relationship between recombination rate and GC content, gene density, and relative distance from chromosome end. The smaller plots on the right show the distribution of the Pearson’s R values from 2000 bootstrap permutations. From left to right, the bottom panel shows the decrease in recombination rate from the end to the center of the chromosome, between-chromosome comparisons of recombination rate and chromosome length, and between-chromosome compairsons of recombination rate and gene density.

## Discussion

Here we provide the first chromosome-scale reference assembly of an intensively-studied Australian bird species, the superb fairy-wren, and the exceptionally diverse family and infraorder to which it belongs. The species has been a focal organism in studying cooperative breeding and sexual selection. This new resource will complement the wealth of behavioural and ecological data, particularly the long-term, multi-decade study of the population in the Australian National Botanical Gardens, Canberra. This reference assembly accompanied by a genetic map and annotation will open up the system to the field of behavioural genomics. This will allow researchers to start searching for genotype-phenotype links behind certain behaviours or behavioural types (Bengston et al., 2018). Researchers can search for candidate genes underlying particular phenotypic traits through gene association studies. This reference assembly would provide information regarding the physical and genetic linkage of multiple genes associated with a particular trait. Researchers can compare gene expression profiles between different phenotypes and this reference can provide insight whether similarities in gene expression are governed by cis- or trans- regulatory regions (Metzger et al., 2016). Researchers will also be able to characterize variation in epigenetic profiles between phenotypes and potentially disentangle the influence of environment or genetics underlying particular traits of interest (Verhoeven, Vonholdt, & Sork, 2016). These are just a few or many possible research avenues of how this reference genome can complement existing research in superb fairy-wrens and its relatives.

The superb fairy-wren genome is particularly important in that it fills a gap in passerine tree as it is the first member of the Meliphagides infraorder to have a reference genome. Not only does it provide a useful resource for species within that clade and the non-passeridan part of the tree but also a useful comparison point in the broader passerine and avian tree for comparative evolution. As part of the early Australasian radiation within oscine passerines, the *Malurus* reference genome anchors the clade which comprises almost half of all avian diversity. This filled gap would allow us to search for shared genomic features which evolved uniquely within the oscine clade. On the other side of the coin, we will also be able to compare how rapidly large scale genomic scale variation has accumulated in oscine passerines, a relatively young radiation, in comparison to the remaining non-passerines.

One initial example of potential for comparative genomics is the evolution of chromosomal rearrangements. Avian genomes have generally been shown to have high synteny (Ellegren, 2010). Previous karyotypic and genome-scale studies, however, revealed that species from certain clades have genomes that have undergone more chromosomal rearrangements than others. Further across birds certain types of rearrangements are more common than others (Ellegren, 2010; O’Connor et al., 2018; Skinner & Griffin, 2012). Although gene synteny along a chromosome might remain high, the chromosome number between bird species have a wide range suggesting that chromosome fusions and fissions might be fairly common (Kapusta & Suh, 2017). The *Malurus* genetic map provides evidence of fusion events between the macrochromosome 2 to microchromosome 19 and macrochromosome 5 to microchromosome 26 (nomenclature based on the *Gallus gallus* reference). More points of comparison would help reveal whether certain chromosomes or particular motifs found in chromosomes might be more prone to rearrangements than others. Furthermore, the dynamic fission and fusion dynamics of the gene-rich microchromosomes may have implications regarding species or clade-specific adaptation (Guerrero & Kirkpatrick, 2014; Hansmann et al., 2009; Wellband, Mérot, & Linnansaari, 2019). Physically creating or breaking linkages between sets of genes will change the local recombination rates. In turn, this would affect the efficiency of selection acting on certain combinations of alleles.

A fine-scale genetic map and recombination landscape is the first step in understanding the causes and consequences of variation in local recombination rates. In particular, a pedigree-based recombination landscape is not as biased by effective population size or selection as a population-based recombination landscape might be. From the existing chromosome-scale genomes, this is only the fifth which is accompanied by a genetic map. In comparison to three of the four available genetic maps, we find that the *Malurus* map is bears more similarity to the *Taeniopygia* map, a distantly related oscine. By contrast, the map for *Ficedula*, which is more closely related to *Taeniopygia*, is more similar to the distantly related *Gallus* (Figure 5). Specifically, recombination deserts on the macrochromosomes, though lower in gene density, could affect patterns of both neutral and adaptive diversity through linked selection (Burri, 2017). This warrants further investigation. We also find variation between male and female maps. Genetic maps from a larger variety of populations and species would be required to start characterizing the variation in this trait and testing hypotheses as to what may be driving this variation between sexes and between species.

Delving into the fine-scale variation within the genome, we found a positive correlation between local recombination rate and GC content, and gene density as well as a stronger and negative relationship with distance from chromosome ends. More broadly we also see a relationship between average recombination rate and chromosome size. These correlations are consistent with previous studies of recombination (Kawakami et al., 2014; Paape et al., 2012). The correlation with GC content is likely due to G-biased gene conversion. Recombination or crossovers during meiosis first require a double-strand break. Although this break is often repaired by using the complementary strand, errors during this repair tend to be biased towards guanine resulting in higher GC content in regions which have high rates of double-strand breaks (Bolívar et al., 2016; Clément & Arndt, 2013; Fullerton, Bernardo Carvalho, & Clark, 2001). Our sliding window analysis shows only a slight positive correlation between gene density and local recombination rate. Selection pressure on local recombination rate would likely depend not only on the density of genes but also the type of genes in a given location and how they are regulated (cis- versus trans-). In the between-chromosome comparison, we found that smaller chromosomes tend to have much higher recombination rate and also tend to have much higher gene density (Axelsson, Webster, Smith, Burt, & Ellegren, 2005). Furthermore, recombination tends to be higher closer to the ends of chromosomes, regardless of the location of the centromeres (Haenel, Laurentino, Roesti, & Berner, 2018). Although we do not yet know the location of centromeres for the *Malurus* genome, this observation stands in this species. While we can only scratch the surface of these comparisons here, understanding the causes and consequences of recombination rate variation is critical in understanding how genome organization influences variation in efficiency of selection across the genome. The recombination rate itself is a unique trait in that it can shift under the influence of selection and yet simultaneously influence the efficacy of selection on various other traits (Comeron Josep M., 2017; Schumer et al., 2018; Wang, Street, Scofield, & Ingvarsson, 2016). Providing these resources from different species would lead us closer to disentangling the complex relationship between the organization of the genome and the evolutionary forces that act on it.

This initial draft of the *Malurus* genome is a well-developed foundation that can be further improved through rapidly developing sequencing technologies and scaffolding methods. As with many of the existing chromosome-scale bird assemblies, not all of the microchromosomes are fully resolved and may still benefit from physical mapping methods such as HiC or BioNano optical mapping (Dudchenko et al., 2017; Jiao et al., 2017; Lehmann et al., 2018). A larger sampling of the pedigree would also provide more informative meioses. That would help in orienting scaffolds already placed in chromosomes. It would also better scaffold the smaller microchromosomes which tend to have higher recombination rate and lower linkage (Fierst, 2015). The capability of long-read sequencing technology has also advanced such that longer fragments can be sequenced. This would help close even more gaps and potentially extend through the telomeres (Kingan et al., 2019; Michael et al., 2018). Highly contiguous, chromosome-scale reference genomes are becoming easily accessible, further expanding our ability to test various hypotheses across. More work can still be done, however, to further refine these assemblies and get closer to the true structure of the genome.

## Acknowledgements

We would like to acknowledge the late Sylvain Foret who guided the design for the sequencing strategy and assembly pipeline of the fairy-wren genome. We would like to thank Kaiman Peng and Angela Higgins from the ACRF Biomolecular Resource Facility at the Australian National University who prepared and sequenced the Illumina mate-pairs and also consulted for the sequencing design. We would like to thank Lutz Froenicke from the DNA Sequencing Facility at UC Davis who guided us through the PacBio sequencing. We would like to thank Andrzej Kilian from the Diversity Arrays Technology who designed and guided us through the sequencing of the pedigree for the linkage map. We would like to thank Takeshi Kawakami and Homa Papoli for advice on linkage mapping and upgrading genome assemblies to chromosome-scale. We would like to thank Ana Catalán for advice on polishing the genome assembly and the BUSCO assessments. Funding was provided by the Australian Research Council (DP150100298), and the office of the DVC (Research) at the Australian National University. Lastly, we would like to thank Mozes P.K. Blom and Claire Peart for comments on the manuscript and various members of the Moritz and Wolf lab for helpful discussions.

## Data Accessibility

*Malurus cyaneus* reference genome: NCBI Accession VKON00000000

NCBI BioSample: SAMN12217597

NCBI BioProject & SRA accession (raw data): PRJNA553115

Genome annotation: TBD

Repeat annotation: TBD

Genetic map & recombination landscape data: TBD

