## Supplemental Material for "Genome of an iconic Australian bird: Chromosome-scale assembly and linkage map of the superb fairy-wren (*Malurus cyaneus*)"

### Supplementary materials

### Supplementary methods

#### Nuclear genome assembly

The following pipeline is organized by the order the steps were completed. The pipeline is not entirely linear but still follows a progression that can be summarized in five stages. A more accurate representation of the pipeline can be found in supplemental figure S2. Each stage was followed by an assessment of contiguity and completeness described in the main text and figure 2.

##### Stage 1: Short-read (Illumina) assembly

###### Illumina data preprocessing

All Illumina reads (shotgun and mate-pairs) were trimmed and filtered using the `trimming.sh` script from *libngs* (https://github.com/sylvainforet/libngs). This script prepares the reads specifically for de novo assembly. The script removes adapter sequences from the standard paired-end libraries and uses *nextclip* (Leggett et al. 2014) to trim the mate-pair libraries. After trimming the Illumina reads proceeded directly to the de novo assembly as well as PacBio error correction.

###### De novo assembly and scaffolding

The Illumina shotgun and mate-pair libraries were assembled and scaffolded using *ALLPATHS-LG* (release 52488) to produce the first assembly stage (Butler et al. 2008). The default settings were used as recommended by the software's developers. Estimates of the insert sizes were provided for the ALLPATHS input. For the shotgun sequencing, the insert sizes were estimated to be 250bp and 500bp with a standard deviation of 100bp. The mate-pair (or "jumping" per ALLPATHS nomenclature) libraries were estimated to be 3500bp, 5500bp and 7500 bp insert size with a standard error of 500bp estimated from the gel excision during library preparation. This stage yielded both a scaffolded and contig assembly.

##### Stage 2: Long-read (PacBio) gap-filling

##### Long-read error correction

Prior to gap filling the PacBio reads needed to be corrected due to its high error rate. Since the coverage of this sequencing is relatively moderate and not high enough to do a self-correction, we used the Illumina reads for correction. We used *LoRDEC* v.0.5 (Salmela and Rivals 2014) to correct the PacBio RSII subreads. This step corrected about half of the reads while the other half remained uncorrected. Both corrected and uncorrected reads were carried through to the gap-filling stage.

```
lordec-correct \
        -i Malurus_PacBio.fq.gz \
        -2 Illumina_reads_1.fastq.gz Illumina_reads_2.fastq.gz … \
        -o Malurus_PacBio_corrected.fastq \
        -k 21 \
        -s 3
```

##### Gap filling

The *ALLPATHS-LG* output included scaffolds with large gaps estimated from the mate-pair libraries. These gaps were filled and some scaffolds were joined using the error-corrected PacBio reads using *PBJelly* (English et al. 2012). Corrected PacBio reads were input as fasta files and the PacBio reads that were not corrected by *LoRDEC* were input as fastq files with the corresponding quality scores. The PBJelly protocol file contained the following *blasr* specification.

```
-minMatch 8 -minPctIdentity 70 -bestn 1 -nCandidates 20 -maxScore -500 -nproc 16 -
noSplitSubreads
```

For the corrected PacBio reads there were no available quality scores. We generated fake quality scores were created using the `fakeQuals.py` script provided in the PBJelly package for those fasta files. The standard PBJelly pipeline was ran with the following modules: setup, mapping, support, extraction, and assembly.

#### Stage 3: Chromosome assembly

#### Putative chromosome assignment

Each scaffold was associated to putative chromosomes based on synteny with the zebra finch genome (v.3.2.4). This step is only used to inform the linkage mapping step when deciding conservative thresholds in LOD scores which determine linkage groups. While other studies use a standard LOD threshold of 3 (i.e. 1000 to 1 chance that linkage observed does not occur by chance), this threshold is quite varied between studies potentially due to variation in recombination or size of the mapping population. The smaller the mapping population, the less power we have to accurately infer linkage. The chromosome assignments are later adjusted using the inferred genetic linkage map and misassembly checks. We used *LASTZ* for the initial chromosome assignment using the following specifications (Harris 2007).

```
lastz ref_chromosome.fa[multiple] Malurus_draft.fa \
        --masking=254 \
        --hspthresh=4500 \
        --gappedthresh=300 \
        --ydrop=15000 \
        --gapped \
        --chain \
        --seed=12of19 \
        --notransition \
        --matchcount=5000 \
        --format=general:name1,start1,end1,length1,name2,start2,end2,strand2 \
        --ambiguous=n \
        --ambiguous=iupac \
        --output=output_name.txt
```

#### Linkage mapping - "Framework map"

The linkage map was generated using a DaRTseq dataseq from a pedigree of superb fairy-wrens (Ren et al. 2015). We followed the *LepMap3* pipeline for the linkage mapping (Rastas 2017). Each module is explained below. This mapping pipeline was carried out specifically so that we can assemble the chromosomes.

***Raw read processing.*** Each DaRTseq raw read library was trimmed to remove Illumina adapter and barcode sequences. We used Trimmomatic (v0.35) with the following specifications with only varying trimming lengths depending on the size of the barcode (Bolger, Lohse, and Usadel 2014).

```
java -jar trimmomatic-0.35.jar SE \
              -phred 33 input.fastq.gz output.fastq.gz \
              -ILLUMINACLIP:TruSeq3-SE:2:30:10 \
              LEADING:3 TRAILING:3 \
              SLIDINGWINDOW:4:15 MINLEN:36
```

***Read alignment.*** The trimmed reads were mapped onto the PacBio gap-filled scaffolds (assembly stage 2) using the default *BWA* specifications (H. Li and Durbin 2009).

```
bwa index Malurus_gapfilled_assembly.fa
bwa mem -t 8 Malurus_gapfilled_assembly.fa trimmed_reads.fq > bwa_align.sam
samtools view -bS bwa_align.sam -o bwa_align.bam
samtools sort bwa_align.bam bwa_align.sorted
samtools index bwa_align.sorted.bam
```

***Preparing LM3 input.*** Since LM3 uses information from genotype likelihoods, we used `samtools mpileup` and scripts provided by LM3 to prepare the input: `pileupParser2.awk` & `pileup2posterior.awk` (Heng Li et al. 2009).

```
samtools mpileup -q 10 -Q 10 -s $(cat sorted_bams)|awk -f pileupParser2.awk|awk -f
pileup2posterior.awk|gzip > post.gz
```

The pedigree was split into 37 full-sibling families with a total of 273 individuals. The posterior file was adjusted into these full-sibling families and the grandparents were included wherever possible. The `post.gz` file and pedigree information were parsed using a python script to produce the `LM_input.post` for the ParentCall2 step.

***ParentCall2.*** This LM3 module adjusts the parent genotype calls using the offspring, grandparents, and half-siblings. This resulted in a total of 35,330 informative markers.

```
java -cp LM3/bin ParentCall2 data=LM3_input.post removeNonInformative=1 halfSibs=1 >
LM3_ParentCall.output
```

***Filtering2.*** The filtering module was run to filter the markers based on high segregation distortion.

```
java -cp LM/bin Filtering2 data=LM3_ParentCall.output dataTolerance=0.01 >
LM3_Filtering.output
```

***SeparateChromosomes2.*** This module separates the markers into linkage groups that should correspond to chromosomes. Multiple iterations of SeparateChromosomes2 was run with changing the LOD score threshold and theta values before the final parameters were run. A LOD score limit of 13 provided the most conservative linkage grouping, where not too many markers from many different putative chromosomes were grouped together in the same linkage group (Figure S3). The LOD score threshold differs considerably from other projects and this may be due to the smaller pedigree sampling.

```
java -cp LM/bin SeparateChromosomes2 data=LM3_Filtering.output \
        lodLimit=13 \
        theta=0.02 \
        numThreads=6 \
        sizeLimit=5 > LM3_SeparateChroms.lod13.theta0.02.size5.output
```

***JoinSingles2All.*** This module adds the unassigned markers into the existing linkage groups using a lower LOD threshold. Trials of LOD thresholds at 3, 4, 5, 7 and 10 were ran to determine a threshold without spurious linkage group assignment. As previously, this is defined by having loci mapped onto scaffolds assigned to many different zebra finch chromosomes grouping in the same linkage group. The higher LOD threshold of 10 was chosen which recovered a total of 11414 loci. These loci were carried through to ordering.

```
java -cp LM/bin JoinSingles2All \
        map=LM3_SeparateChroms.lod13.theta0.02.size5.output \
        data= LM3_Filtering.output \
        lodLimit=10 \
        iterate=1 > LM3_JoinSingles.lod10.output
```

***OrderingMarkers2.*** This module orders markers within each linkage group to reflect the linkage map. This module required multiple iterations and some manual curation as it would determine the ordering and orientation of each contig within a chromosome. The ordering was run separately for each putative chromosome. This initial command was run 10 separate times with

```
java -cp LM/bin OrderMarkers2 \
        data=LM3_Filtering.out \
        map=LM3_JoinSingles.lod10.output \
        numThreads=6 \
        sexAveraged=1 \
        chromosome=1 \ #[whichever chromosome was being run]#
        > LM3_OrderMarkers_chr1.trialX.txt
```

6 iterations within each run.

This process was repeated until the map did not seem to have any additional spurious markers. This initial map yielded 26 linkage groups with 11,219 loci. We refer to this as the “framework map” from which we will build a forced map that will be used for the scaffolding.

#### Chimeric scaffold detection and correction

Putatively misassembled scaffolds were identified using multiple criteria to prevent misidentification of true chromosomal fusions or translocations.

(1) Putative chimeric scaffolds were first identified using the linkage map. Scaffolds where one end contained a series of consecutive SNPs that belonged to a linkage group different to the rest of the SNPs on the scaffold were flagged as chimeric. This criteria found 6 putatively chimeric scaffolds.

(2) All scaffolds were also queried against the zebra finch and flycatcher chromosomes. Scaffolds that contained multiple regions larger than 15Kb that matched different chromosomes in both assemblies were flagged as chimeric. This found 14 additional putatively chimeric scaffolds.

The scaffolds that were identified using the linkage map were also recovered in the LASTZ query. This initial identification allowed us to narrow down the scaffold breakpoints to ~100Kb window. The LASTZ parameters were the as described in the 'Putative Chromosome assignment' section.

To narrow down the breakpoint further, we mapped the three mate-pair libraries onto the draft genome assembly using BWA-mem. We used the insert size information provided by the ALLPATHs assembly output as the BWA-mem insert size distribution. We then extracted the number of reads with mates mapping onto a different scaffold within 10kb non-overlapping windows across each scaffold. Within the initial 100kb breakpoint window, we identified regions with unusually high number of matching mate reads mapping to other scaffolds. These regions likely contain repetitive elements which may have been responsible for the assembly of these chimeric scaffolds in the first place. These regions were also visually inspected using IGV viewer (Robinson, Thorvaldsdóttir, and Winckler 2011). The breakpoint was narrowed down to 10-20kb, them that region was completely removed and the chimeric scaffold was split.

```
bwa mem —t 4 —p reference_db_prefix \
        mate_pair_library.fq.gz \
        —I 2331[,494] > output.sam

## 3.5kb insert library —I 2331[,494]
## 5.5kb insert library —I 3944[,583]
## 7.7kb insert library —I 5968[,1620]
```

The total length of the affected sequence was ~86Mb. Eight larger scaffolds were split into 17 smaller scaffolds to be used for the placing and orienting.

#### Linkage mapping - "Forced map"

After we were fairly confident about splitting the chimeric scaffolds, we generated a forced map which uses the information from the physical map to assign more SNPs into the map. First we determined which scaffold was assigned to one of the 26 linkage groups, then we determined which SNPs were mapped onto those scaffolds but not included in the framework map. We forced all of the SNPs within a scaffold into the linkage group which the scaffold belonged to in the framework map. This resulted in a large 32,708 SNPs (out of the 35,276 SNPs) to be assigned in one of the 26 linkage groups.

***JoinSingles2All***. To maximize the numbers of scaffolds assigned to linkage groups, we used this large forced map and attempted to join any of the remaining unmapped SNPs into one of the existing groups. We ran the JoinSingles2All module with a LOD score cut off 7. This step resulted in 33,098 SNPs assigned to the 26 linkage groups.

***SeparateChromosomes2***. Considering that this forced map might have SNPs which are not informative or have unusual heritability, we filtered the SNPs within each linkage group using the SeparateChromosomes2 module. For each linkage group, we made it such that the pool of SNPs are only those that mapped to associated scaffolds. We then ran the SeparateChromosomes2 with a LODlimit=3 within each linkage group and recovered the largest linkage group. This results in a forced map where all SNPs within a linkage group should be associated with at least a LOD score of 3. This results in 26,157 SNPs in the 26 linkage groups

***OrderingMarkers2.*** Finally, we did a last iterative round of OrderingMarkers2 to get the order within a linkage group. We ran this as extensively as described for the framework map and also carried out manual curation. This manual curation refers to removal of SNPs that seem to have been placed incorrectly as indicated by high genetic distances and constantly being placed at the end of the marker order. This stage also dropped one of the linkage groups as it was only associated to one scaffold which is large but not large enough to constitute a chromosome. The final map which we used to build the chromosome scale assembly contained 25,078 SNPs in 25 linkage groups.

##### Anchoring and orienting scaffolds into chromosome superscaffolds

After the putative chimeric scaffolds were split, the final linkage map was used to place and orient the scaffolds into chromosome-scale super scaffolds. ALLMAPs was used to build 25 superscaffolds (Tang et al. 2015). For all of the chromosomes, both the male and female linkage maps were merged into a single bed file. For the Z chromosome, only the male linkage map was used.

```
python -m jcvi.assembly.allmaps merge malurus_male.csv malurus_female.csv -o
malurus_merged.bed
python -m jcvi.assembly.allmaps path malurus_merged.bed Malurus_draft.fa
```

ALLMAPs places a standard 100bp interscaffold gap size following the Genbank convention. This scaffolded draft was then subjected to another round of PacBio gap filling.

#### Stage 4: Chromosome gap-filling

##### Gap filling

The parameters used for this second round of gap filling were identical to the parameters used for the initial round of gap filling described above. The rationale for running a second round of gap filling is that approximately 5Mb of new gaps were introduced during the upgrade from scaffolds to superscaffolds. Additionally, the new placement of scaffolds may provide the necessary backbone for new mapping locations for the PacBio reads.

#### Stage 5: Genome polishing

##### Pilon polishing

The final stage is the assembly polishing. We used Pilon (v.1.22) to correct the incorrect bases, small indels, small gaps, and local misassemblies in the final gap-filled superscaffolds (Walker et al. 2014). First, we mapped all the cleaned Illumina reads using identical parameters used in the chimeric scaffold correction step. Each of the libraries with different insert sizes was mapped separately. Second, we ran Pilon on small chunks of the genome to accommodate the large memory requirements. The polishing was done in chunks ranging from 2.5Mb - 10Mb.

```
java -Xmx400G -jar $PILON_HOME/pilon.jar \
        --genome ${genome_path}/${genome} \
        --frags ${align_path}/Malurus_cyaneus_250bp_paired.sorted.bam \
        --frags ${align_path}/Malurus_cyaneus_500bp_paired.sorted.bam \
        --jumps ${align_path}/Malurus_cyaneus_2kb_matepair.sorted.bam \
        --jumps ${align_path}/Malurus_cyaneus_4kb_matepair.sorted.bam \
        --jumps ${align_path}/Malurus_cyaneus_6kb_matepair.sorted.bam \
        --unpaired ${align_path}/Malurus_cyaneus_250bp_single.sorted.bam \
        --unpaired ${align_path}/Malurus_cyaneus_500bp_single.sorted.bam \
        --threads 16 \
        --diploid \
        --changes \
        --vcf \
        --targets chunk_N \
        --output chunk_N_out \
        --outdir outdirectory
```

##### Final chromosome assignment

The assembled chromosomes were assigned their associated number when the scaffolds were queried against the zebra finch genome assembly prior to the genetic linkage mapping stage ("Initial assignment" section). Chromosomes that fused were assigned the number of the larger chromosome. This resulted in a total of 25 chromosome assemblies.

The remaining scaffolds were queried against the chromosome-scale assemblies of the *Ficedula albicollis* (v1.5), *Taeniopygia guttata* (v.3.2.4), *Passer domesticus* (v1), and *Parus major* (v1) using the same LASTZ specifications previously. If a scaffold is found to be associated with the same chromosome of at least two of the four genomes, it was assigned to that chromosome but written out to the "unplaced scaffolds" files. Chromosomes for which we did not have an associated linkage group (likely due to the low number of loci or individuals) but had recovered large scaffolds were retained. The largest scaffold was used to represent those chromosomes (Chromoso...).

Lastly, a few scaffolds were assigned to the W chromosome though unplaced. Since the sequenced individual was female, some of the scaffolds are likely from chromosome W but has consistently been proven difficult to assemble. We assigned scaffolds to the W chromosome using the genotype information from the pedigree mapping population. If SNPs were consistently only genotyped in females and were only homozygous, these were good candidates for W-linked SNPs.

#### Assembly quality assessment

The quality of the assemblies was assessed after each major stage. Other chromosome-scale genomes were also assessed in the same manner so that we have a comparison point. Standard genome assembly metrics regarding sizes of scaffolds, contigs, and gaps were calculated using a perl script in the Assemblathon 2 analysis (https://github.com/ucdavis-bioinformatics/assemblathon2-analysis). This assessment is to observe how each step improves the contiguity of the assembly as well as how many gaps were filled during the process.

The other method to assess genome quality is through the standard BUSCO search (v3; (Simão et al. 2015)). This method assesses completeness by searching the assembly for universal single-copy orthologs. We used the Aves (odb9) database that contains 4915 BUSCO genes. Recovery of intact single-copy orthologous genes is a proxy for the completeness of the genome assembly. Improvements in the number of BUSCO genes recovered also tends to reflect the improvement in assembly quality in each stage.

### Mitochondrial genome assembly

Since the mitochondrial genome as not recovered when the ALLPATHS assembly was queried using BLAST, we assembled the mitochondrial genome separately using MITObim (Hahn, Bachmann, and Chevreux 2013). A complete mitochondrial genome assembly of the congeneric red-backed fairy-wren (*Malurus melanocephalus*) from GenBank (NC_024873) was used for the initial seeding and recovery of mitochondrial reads. Multiple trials were run using different

MITObim configurations and using either the 150bp PE HiSeq2500 reads or the 300bp PE MiSeq reads. The 300bp PE MiSeq reads consistently yielded assemblies that were significantly longer than expected likely due to assembling through NUMTs in the nuclear genome. After the trials, the MITObim assembly using the 150bp PE with the 'mapping' mode yielded the best genome after 5 iterations.

After the MITObim assembly, we assessed the quality of the genome by mapping the 150bp PE

```
perl MITObim.pl \
	-end 10 \
	-sample malurus_cyaneus \
	-ref malurus_melanocephalus \
	-readpool malurus_cyaneus_150bp_PE.fastq.gz \
	--quick malurus_melanocephalus_ref.fa \
	--pair \
	--mira_path /path/to/mira/executables
```

reads onto the assembly using BWA-mem. We then checked for approximately even coverage

```
samtools mpileup -Ou -f mitobim_genome.fa bwa_alignment.sorted.bam | bcftools call -mv -Oz -
o malurus_mitogenome.vcf.gz

tabix malurus_mitogenome.vcf.gz

cat mitobim_genome.fa | bcftools consensus malurus_mitogenome.vcf.gz >
malurus_mitogenome.consensus.fa
```

using the IGV Genome Browser. Any large spikes in coverage may indicate NUMTs being misassembled. No large spikes in coverage was recovered in the final alignment. The BWA alignment was also using to correct any erroneous bases from the MITObim assembly by taking the most common base of the mapped reads. We used samtools (1.3.1), bcftools (1.2), and tabix (1.2.1) (Heng Li et al. 2009; Heng Li 2011).

The mitochondrial genome underwent an initial annotation using the MITOS Webserver to identify the location the genes and tRNAs (Bernt et al. 2013).

#### Linkage and recombination maps

#### Genetic linkage map

We ran the LM3 pipeline again to generate the final linkage and recombination maps. The DArTseq reads were remapped to the whole chromosomes in the final genome assembly as the additional gap filling and polishing could have fixed sites so that more SNPs could be used for the map. We did not include the SNPs in the unplaced or unknown scaffolds. This resulted in 35,349 SNPs mapping onto whole chromosomes.

We ran most of the pipeline as we did previously but followed the steps in the **SeparateChromosomes2** subsection in the forced map so that within each chromosome, we only estimated the linkage between SNPs that are associated with a LOD score of 3. Prior to estimating the map distances, we ordered the SNPs based on the physical order along the chromosome. Any mismatches in the genetic order and physical order would result in unusual jumps in the linkage and was filtered out. The mismatches can be due to error from a small mapping population, differences between subspecies from which the genetic map and physical map are sequenced, or polymorphic structural variation within the mapping population.

We estimated the sex-specific and sex-averaged maps for all 24 autosomes and estimated the male-specific map from the Z chromosome.

#### Recombination map

Estimating the recombination map involves translating the centimorgan measure in the genetic map into centimorgans per Megabase for local, sliding-window recombination rates. We did this by using the Marey map configuration of the genetic map where the Y-axis is the genetic map and the X-axis is the physical map (Chakravarti 1991). We then did a Loess (locally weighted scatterplot) smoothing so that we have a smoothed curve determined by sliding windows of fixed numbers of SNPs. This ensures there is no inherent bias in regions where we have high SNP density. We used the loess function in R with a span of 0.2 to fit a curve along the Marey map for each chromosome. To then estimate the recombination rate (cM/Mb) in any arbitrary sized sliding window, we used the genetic distance as predicted by this curve within the window (cM) and divided it by size of the window in units of centimorgans.

The resulting recombination maps can be found in figure S8. We also correlated the sliding window recombination rates (200kb, nonoverlapping windows) with various genomic features to test correlations within the genome. This is discussed in detail in the main text and the correlation figure 5.

#### Repeat annotation

We ran a simple repeat annotation to roughly assess the repeat content of the genome. We ran RepeatModeler (v.1.0.8) for de novo repeat identification in the *Malurus* genome assembly (Smit et al. 2013-2015). We then used the resulting de novo identified repeats from RepeatModeler and repeat libraries from other bird species to find and mask the repeats in the final assembly.  RepeatMasker were run (Smit et al. 2008-2015). We used the database containing libraries from the zebra finch to train the model to find more repeats in the fairy-wren genome. First, we ran RepeatModeler on the polished genome assembly with the following standard specifications:

```
BuildDatabase —name Malurus_db Malurus_polished_genome.fasta
RepeatModeler —pa 4 —database Malurus_db
```

### Supplementary figures

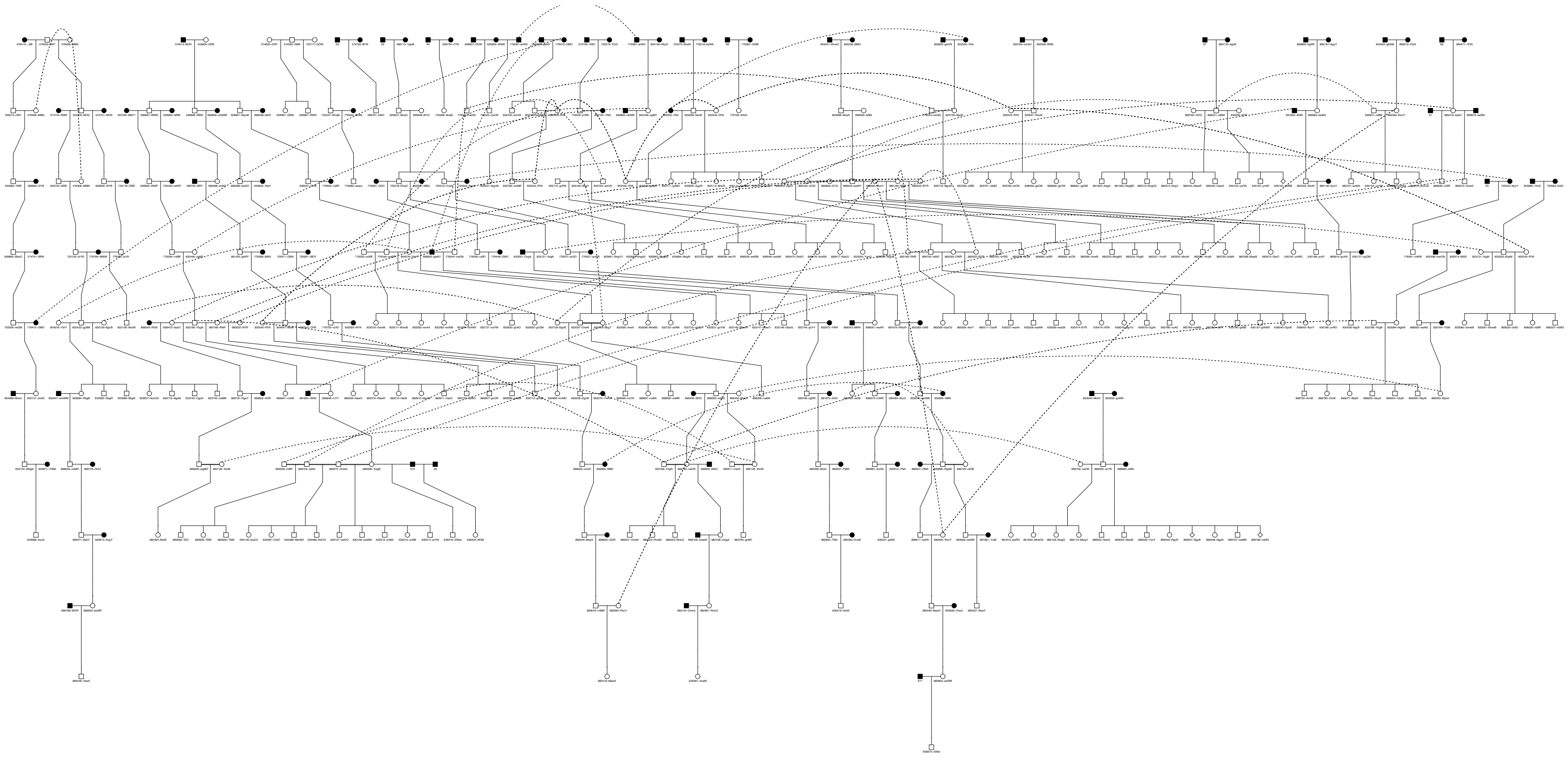

**Figure S1**. Pedigree of the wild population used for the linkage map. Individuals in black represent those unsampled for sequencing. [illegible]

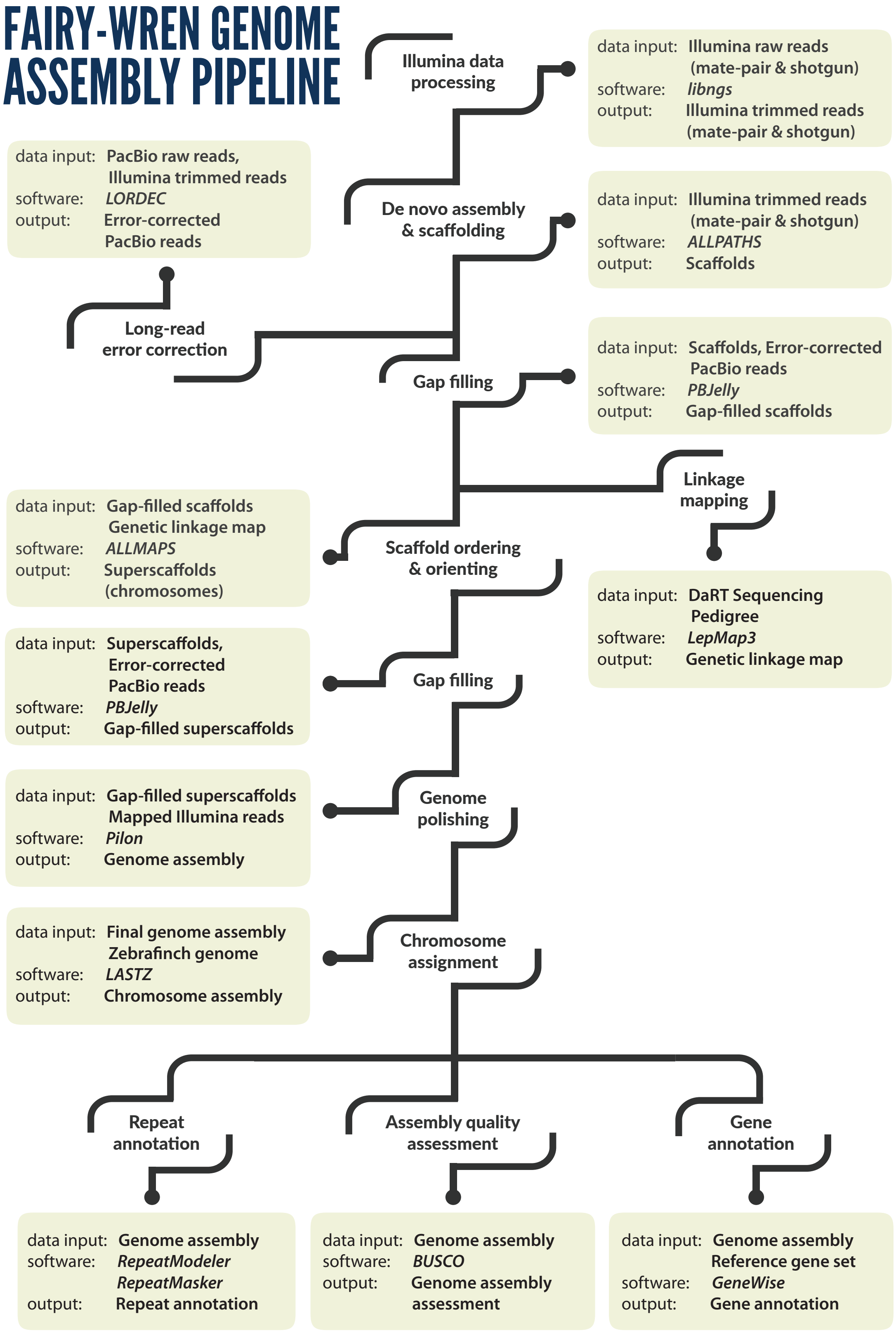

**Figure S2.** Summary of the nuclear genome assembly pipeline.

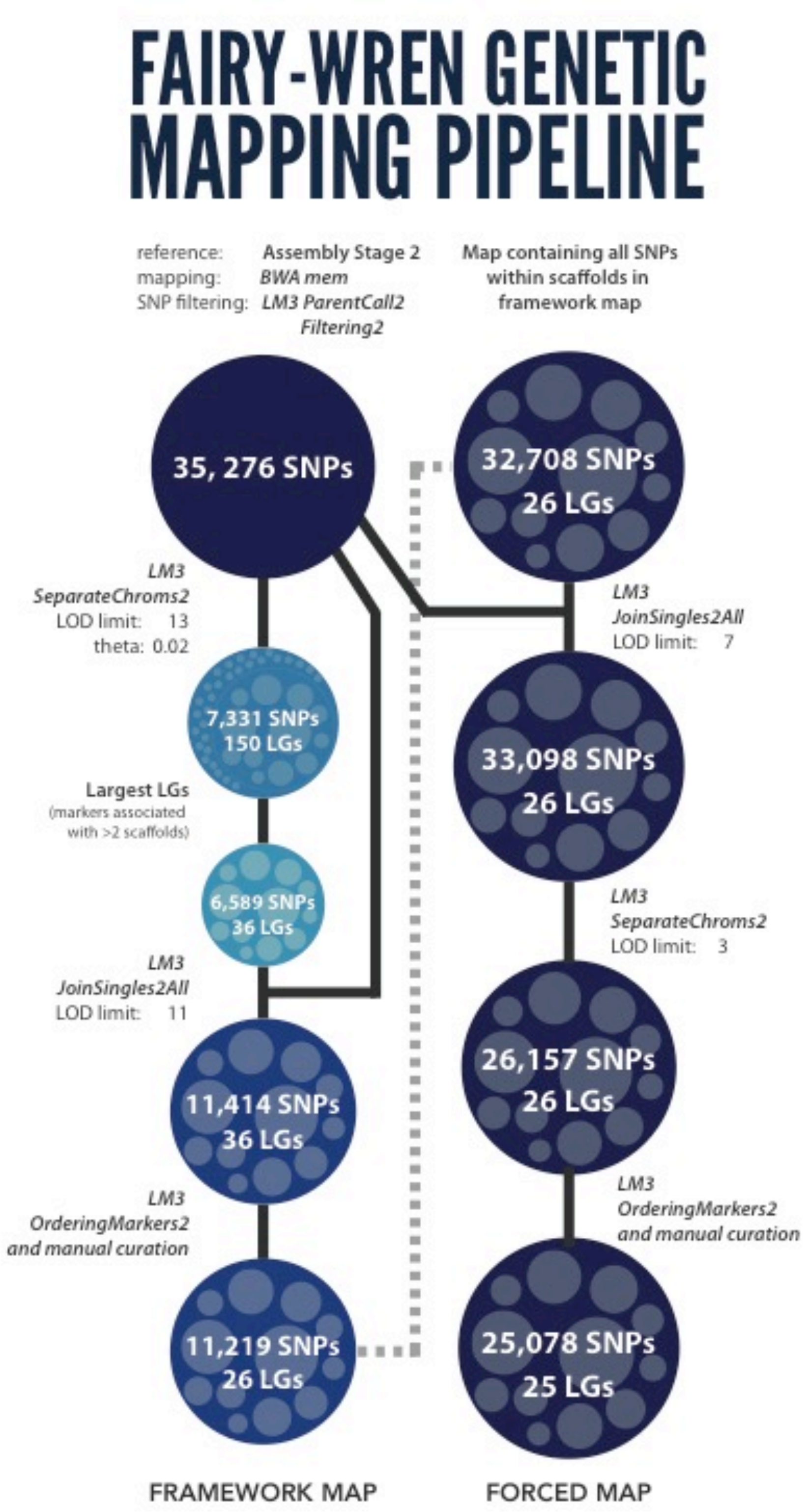

**Figure S3.** Summary of the linkage mapping pipeline through the LepMap3 mapping software. The framework map was generated conservatively to serve as the backbone for the forced map. The final forced map was used to anchor and orient scaffolds into chromosomes.

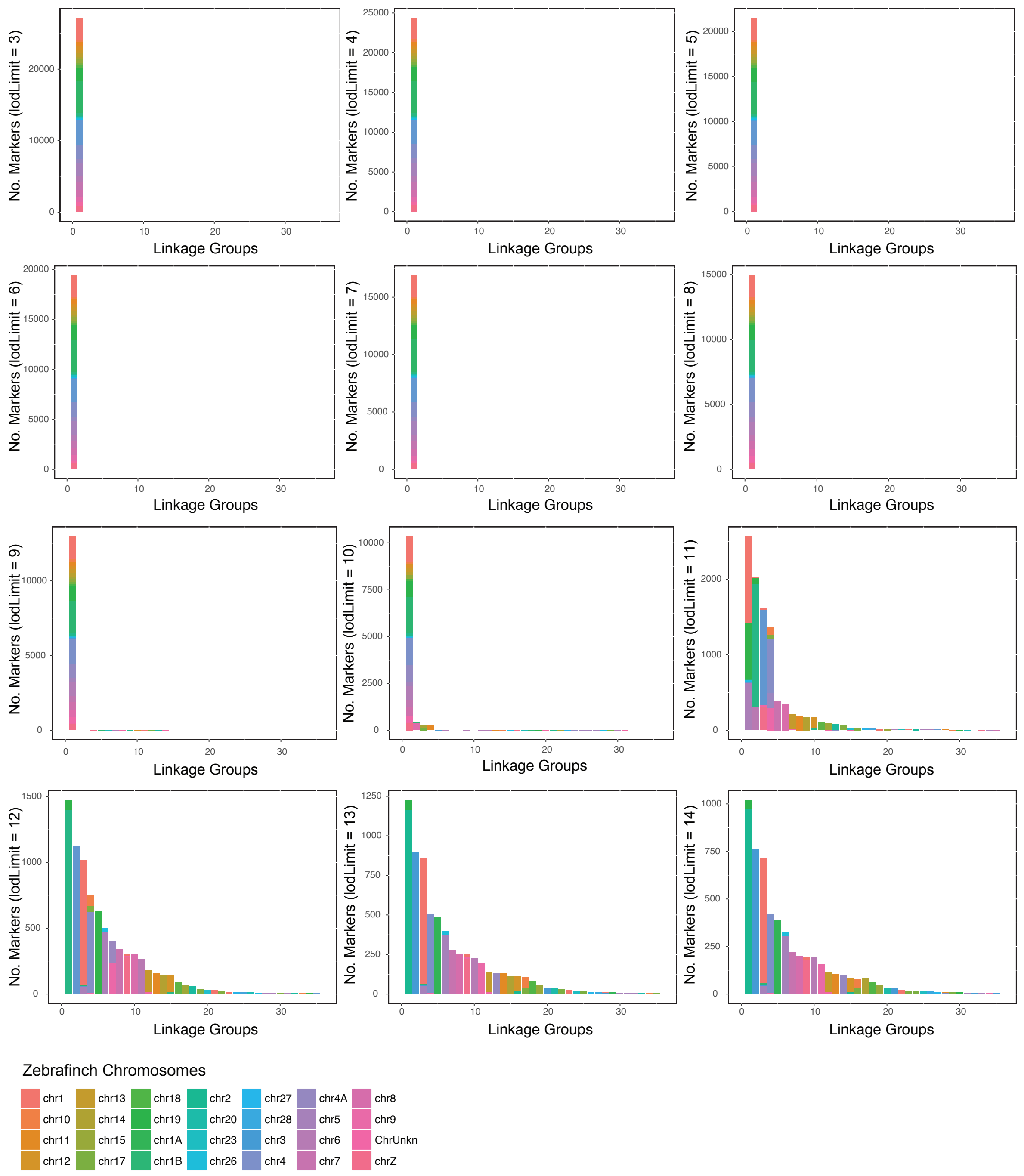

**Figure S4.** Trials of different LOD score cutoffs to determine the most conservative threshold. The scaffolds were first mapped to the zebrafinch genome assembly (v.3.2.4) to have a roughly informed cut off. A threshold below LOD=11 tend to group most loci in a single linkage group. The threshold at LOD=13 is the lowest where the loci tend to be placed in a linkage group that corresponds to homologous zebrafinch chromosomes.

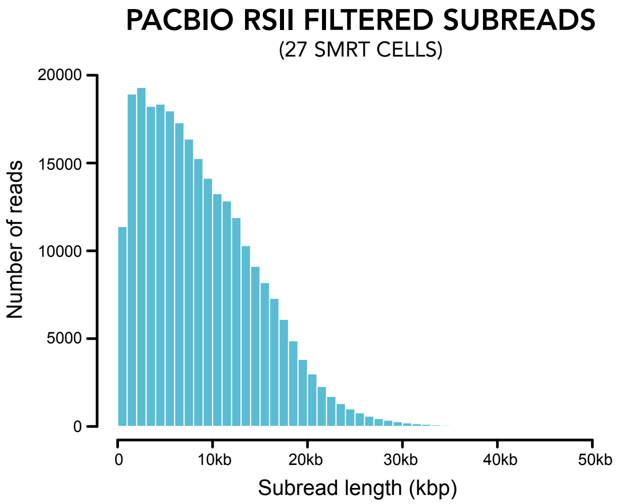

**Figure S5.** Histogram of the PacBio RSII filtered subread lengths. In total we sequenced 2,676,850 subreads, corresponding to ~22x of the superb fairy-wren genome.

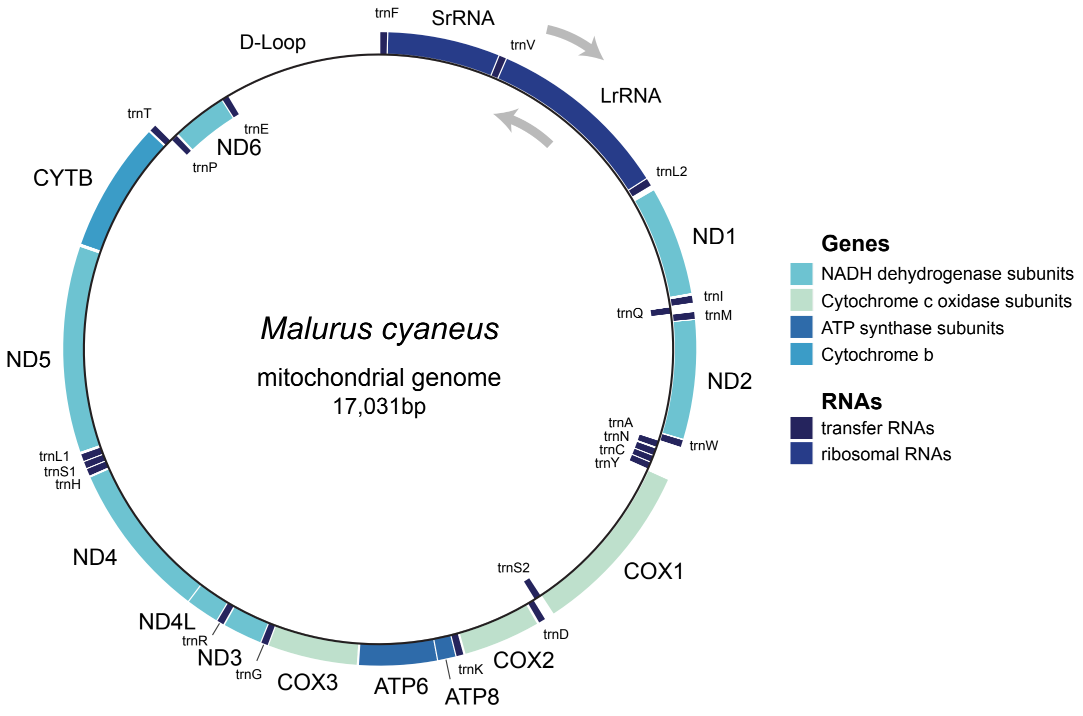

**Figure S6.** Mitochondrial genome assembly and annotation. The mitochondrial genome was assembled using MITObim and annotated using the MITOs Webserver.

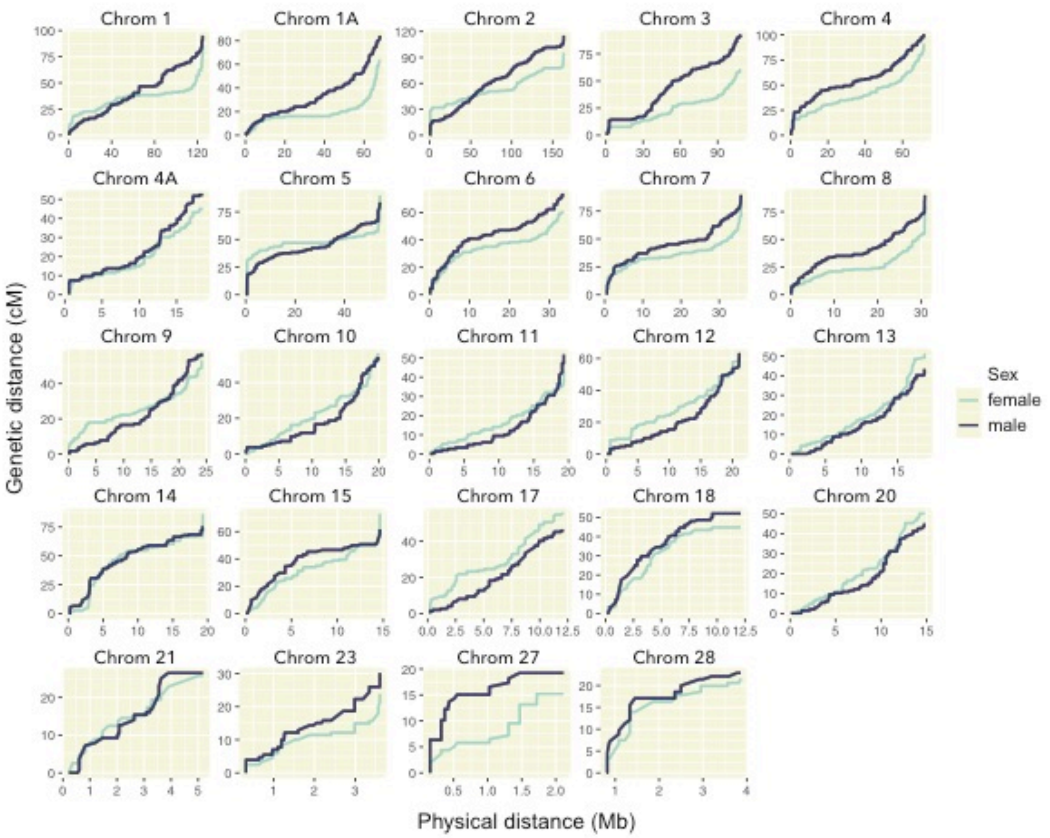

**Figure S7.** Sex-specific Marey maps of the superb fairy-wren autosomes.

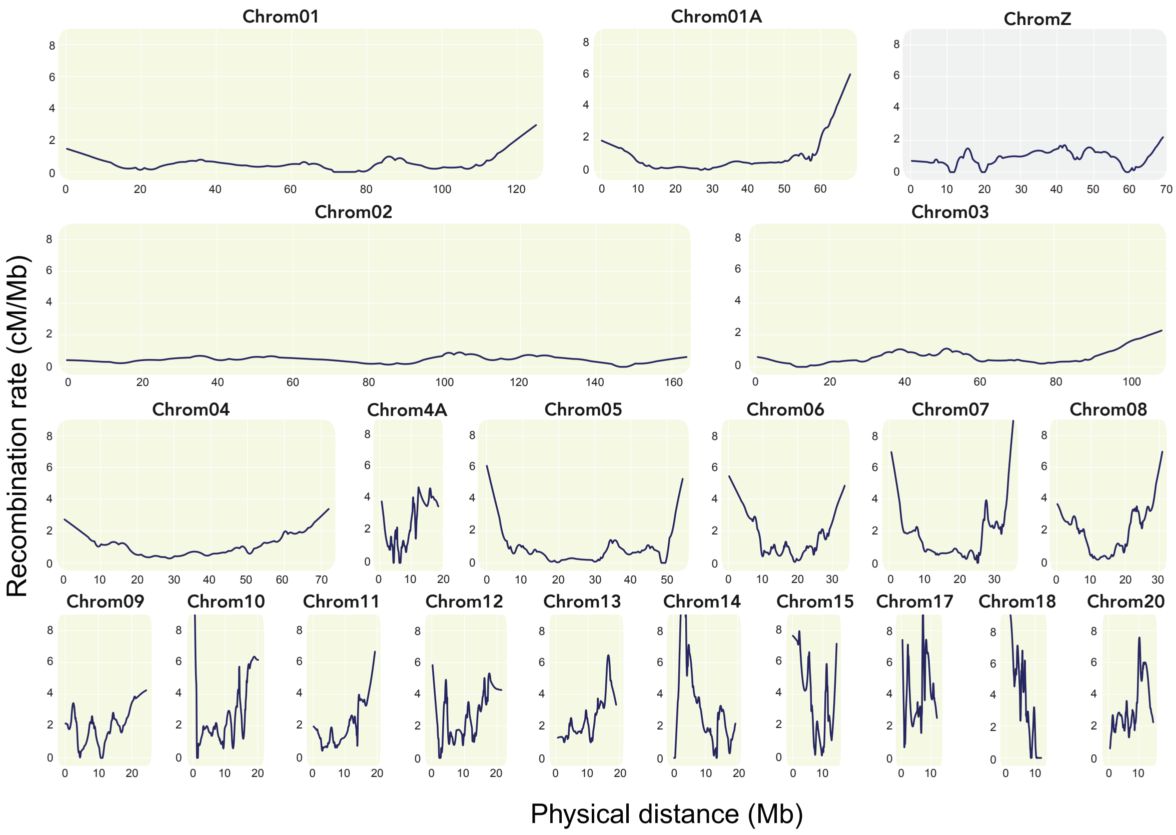

**Figure S8.** Recombination rate landscapes across the chromosomes.

### Supplementary tables

Table S1. Summary table of sequencing strategy and resulting raw data.

| Lib. Type | Seq. Platform | Read lengths | Insert size | Seq. effort | Number of reads | Coverage |
|---|---|---|---|---|---|---|
| shotgun | Illumina HiSeq2500 | 100bp PE | 200-400bp | ½ lane | 91 417 772 pairs | 18.93x |
| mate-pair | Illumina HiSeq2500 | 100bp PE | 2-3kbp | ⅙ lane | 12 287 646 pairs | 1.46x |
| mate-pair | Illumina HiSeq2500 | 100bp PE | 4-5kbp | ⅙ lane | 15 656 984 pairs | 1.86x |
| mate-pair | Illumina HiSeq2500 | 100bp PE | 7-8kbp | ⅙ lane | 9 326 889 pairs | 1.11x |
| shotgun | Illumina MiSeq | 300bp PE | 450-550bp | 1 lane | 6 303 745 pairs | 4.81x |
| long-read | PacBio RSII | See Fig. S1 | See Fig. S1 | 27 CELLs | 2 676 850 subreads | 23x |

Table S2. RepeatMasker summary table. Of the total genome length of 1.07Gb, 85.21Mb (7.90%) was masked. The database used contained repeats derived from *Taeniopygia guttata*, *Ficedula albicollis, Corvus cornix*, and de novo *Malurus cyaneus*.

| | Number of Elements | Length Occupied | | Percentage of Sequence |
|---|---|---|---|---|
| SINEs: | 7080 | 1143019 | bp | 0.11% |
| ALUs | 0 | 0 | bp | 0.00% |
| MIRs | 2131 | 226185 | bp | 0.02% |
| LINEs: | 123748 | 35048152 | bp | 3.25% |
| LINE1 | 26 | 2796 | bp | 0.00% |
| LINE2 | 0 | 0 | bp | 0.00% |
| L3/CR1 | 123645 | 35030170 | bp | 3.25% |
| LTR elements: | 46441 | 24305581 | bp | 2.25% |
| ERVL | 29895 | 16034660 | bp | 1.49% |
| ERVL-MaLRs | 0 | 0 | bp | 0.00% |
| ERV_classI | 9811 | 5088530 | bp | 0.47% |
| ERV_classII | 2259 | 1613144 | bp | 0.15% |
| DNA elements: | 5505 | 828729 | bp | 0.08% |
| hAT-Charlie | 149 | 52630 | bp | 0.00% |
| TcMar-Tigger | 173 | 29583 | bp | 0.00% |
| Unclassified | 21827 | 70310963 | bp | 0.65% |
| Total interspersed repeats: | | 68354544 | bp | 6.34% |
| Small RNA: | 1684 | 526507 | bp | 0.05% |
| Satellites: | 1508 | 398457 | bp | 0.04% |
| Simple repeats: | 252826 | 13132276 | bp | 1.22% |
| Low complexity: | 51834 | 3233359 | bp | 0.30% |
